# Taking a BREATH (Bayesian Reconstruction and Evolutionary Analysis of Transmission Histories) to simultaneously infer phylogenetic and transmission trees for partially sampled outbreaks

**DOI:** 10.1101/2024.07.11.603095

**Authors:** Caroline Colijn, Matthew Hall, Remco Bouckaert

## Abstract

We introduce and apply Bayesian Reconstruction and Evolutionary Analysis of Transmission Histories (BREATH), a method to simultaneously construct phylogenetic trees and transmission trees using sequence data for a host-to-host outbreak. BREATH’s transmission process that accounts for a flexible natural history of infection (including a latent period if desired) and a separate process for sampling. It allows for unsampled individuals and within-host evolution. It also accounts for the fact that an outbreak may still be ongoing at the time of analysis, using right truncation adjustment. We perform a simulation study to verify our implementation and explore sensitivity to unknown or misspecified parameters. We then apply BREATH to a previously-described 13-year outbreak of tuberculosis. We find that using a transmission process to inform the phylogenetic reconstruction results in better resolution of the phylogeny (in topology, branch length and tree height) and a more precise estimate of the time of origin of the outbreak. Considerable uncertainty remains about transmission events in the outbreak, but our reconstructed transmission network resolves two major waves of transmission consistent with the previously-described epidemiology, estimates the numbers of unsampled individuals, and describes some high-probability transmission pairs. An open source implementation of BREATH is available from https://github.com/rbouckaert/BREATH as the `BREATH` package to BEAST 2.

## 1. Introduction

Genomic data are increasingly used in infectious diseases surveillance for both human and animal populations. Declines in the cost of sequencing, and the utility of pathogen sequence data to support epidemiological investigations, have made them appealing for a range of applications. In particular, pathogen genomes can be used to refine knowledge of transmission clusters, to identify links between previously unlinked epidemiological clusters, to characterize the timing of transmission, and to identify likely transmission pairs or refute putative ones [25, 33]. These can, in turn, yield information about the host- and strain-related risk factors for onward transmission [34].

There are substantial challenges involved in inferring full transmission trees from genomes. Multiple pairs of individuals may have very closely-related or even identical pathogen genomes [13, 20]. Hosts may also harbour substantial pathogen diversity, particularly for chronic infections such as tuberculosis. These both complicate the connection between pathogen sequence relatedness and whether one host infected another. The timing of symptoms or case detection is not very informative about the infection time for some diseases, including chronic infections such as tuberculosis [15]. The existence of unobserved, or observed but unsequenced, hosts is likely in most settings and complicates the relationship between pathogen sequences and who infected whom. Despite these challenges, pathogen sequence data are a potentially valuable source of information, as sequences encode variation that is informative about ancestry.

A range of methods have been developed to reconstruct who infected whom using sequence data [29, 32]. These methods include (to name a few) outbreaker2 [19], which integrates epidemiological and genetic data and accounts for unsampled hosts, but does not account for within-host evolution or the shared evolution of the sequences (e.g. through a phylogenetic tree); TransPhylo [17, 26, 35], which accounts for within-host evolution and unsampled hosts but requires a fixed timed phylogenetic tree as an input; and BEASTLIER [12] and phybreak [18], which jointly estimate the phylogenetic tree and transmission tree, accounting for within-host evolution, but do not allow for unsampled hosts.

Phylogenetic trees are a key tool to analyze sequence data, because they are constructed with molecular evolutionary models that allow for rate heterogeneity (both between types of base substitutions and genomic positions), molecular clock estimation, consideration of genome content, and other complexities [10, 6, 14, 21]. They also describe the shared ancestry of the set of taxa, and this shared ancestry embeds relationships among the possible transmission pairs that relate to the possible transmission trees. Accordingly, it is an advantage to use phylogeneties in reconstructing transmission, compared to, for example, pairwise distances and treating pairs as independant. However, at the short time scales of person-to-person transmission, we likely do not have a large number of genetic polymorphisms with which to reconstruct a single phylogenetic tree with high confidence [20]. For this reason, TransPhylo has been adapted to allow for simultaneous inference on multiple input phylogenies to account for phylogenetic uncertainty [24]. This is time consuming, and the sample of phylogenies for which it is feasible to re-run an MCMC inference is unlikely to capture the phylogenetic uncertainty very well. In addition, since the phylogeny is uncertain, it is helpful to inform phylogenetic reconstruction with prior knowledge about the process that generated the data, namely a transmission process. This cannot be done if the phylogenetic trees are fixed prior to transmission analysis.

Here we introduce a method called Bayesian Reconstruction and Evolutionary Analysis of Transmission Histories (BREATH). Its primary novelties are that (1) it jointly estimates transmission events and the timed phylogenetic tree, allowing for unsampled hosts and within-host evolution and (2) it uses a very flexible model for timing which can account for real-time outbreaks in which there is right truncation. The joint inference of phylogeny and transmission allows the phylogenetic tree to be informed by knowledge of the transmission process (e.g. the likely time between infection and infecting others; likely time from infection to sampling; the fact that the pathogen is spread host-to-host, or premises-to-premises). We use a likelihood based on intensity functions and we account for the fact that only individuals who are “ancestral to the sample” can be known to the model. We implemented BREATH as a package of the BEAST 2 software platform [22], opening access to the wide variety of models and methods already implemented in BEAST 2 and its packages, and to state-of-the-art phylogenetic reconstruction.

## 2. Methods

In overview, BREATH is a method to construct transmission trees and phylogenetic trees at the same time, using a Bayesian approach to account for uncertainty in both. It assumes host-to-host transmission with a given (perhaps high-variance) generation time distribution and distribution of time from infection to sampling. It accounts for within-host evolution, allowing hosts to have more than one pathogen lineage at a time. BREATH accounts for unsampled infections, and so does not require full sampling.

### 2.1. Mapping transmission on to a phylogeny

BREATH annotates a timed phylogeny with a transmission tree, using the same kind of “colouring” approach introduced previously in both TransPhylo and BEASTLIER [9, 17, 12]. In these works, each point on the phylogeny was associated with a specific individual host, whether sampled or unsampled. Note that a “host” may in some applications be an infected location such as a farm rather than an individual human or animal. Here, we also model transmission chains of unsampled hosts that can occur on a branch of the phylogeny. As a result we have two classes of unsampled hosts: “individual” unsampled hosts (IUHs) and members of chains of unsampled hosts. These chains have a duration, making up a subinterval of the branch that they take place on; we keep track of the number of hosts in a chain. Figure 1 illustrates the colouring, sampled an unsampled hosts, chains of unsampled transmission, and shows a phylogeny and the corresponding transmission tree.

**Figure 1:**
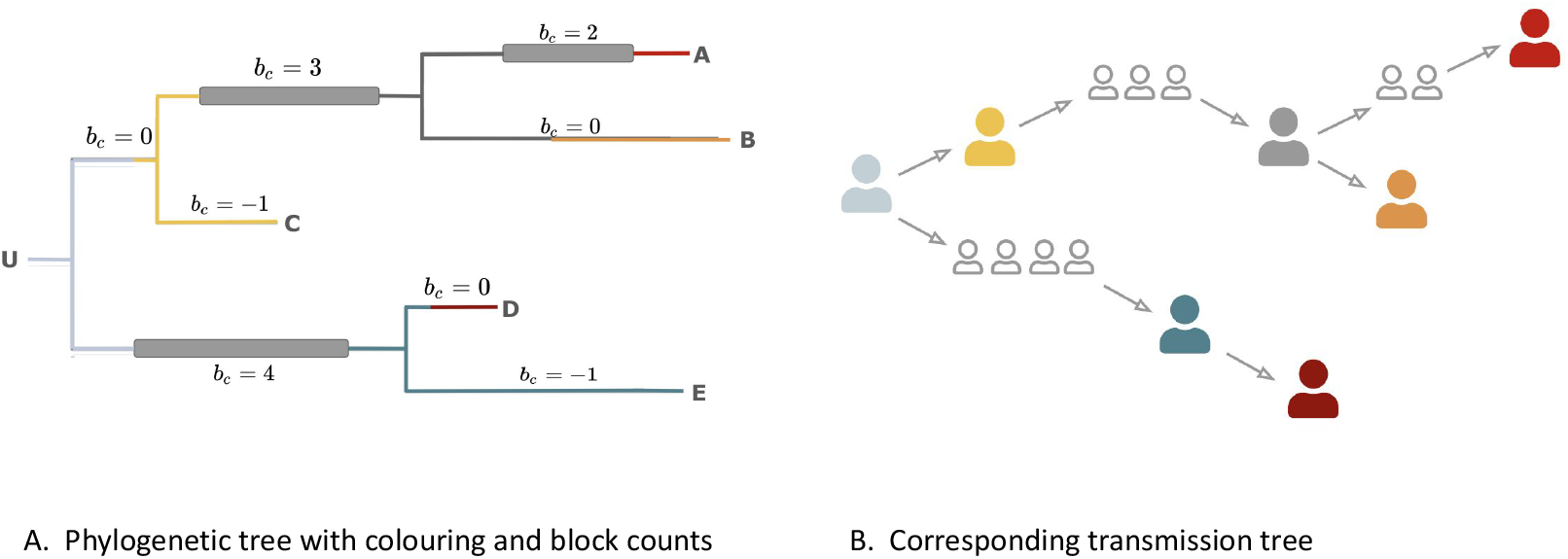
An example of a coloured phylogeny (A) and corresponding transmission tree (B). The transmission process starts with an individual unsampled host U in whom the root node resides. U infects C, who is sampled. C hosts the yellow portion of the tree in panel A and is yellow in panel B. U also starts a chain of unsampled transmission of length 4. This chain is represented by the block labelled *b*_*c*_ = 4 in panel A and the chain of 4 individuals in panel B. C starts a chain of unsampled transmission of length 3. The chain infects an IUH, who infects B (orange in both panels A and B). B is sampled. The IUH also infects a chain of length 2, which infects A (red in both panels), who is sampled. In the lower clade, the chain of length 4 infects E (blue), who infects D (dark red). They are both sampled. The “block counts” (see *Implementation*, Section 2.4) are labelled.

Our model allows for within-host evolution in that a host can harbour multiple pathogen lineages at the same time. The within-host evolution allows for transmission events to be distinct from branching events, and greatly increases the number of transmission trees that are consistent with a given phylogeny. For example, in Figure 1, the grey individual unsampled host infects case B (orange) and a chain of length 2 that infects A (red). B receives a different lineage than the first individual in that chain, and the most recent common ancestor of those two transmitted lineages existed some time prior to either transmission event. Without the within-host diversity, transmission events would occur only at phylogenetic nodes, and thus the TMRCA of the transmitted lineages would have to take place at the same time as the first of the transmissions. This is particularly appropriate in a chronic infection where considerable diversity can accrue within hosts. Within-host evolution affects the relationship between phylogenetic trees and transmission trees, even if it is not measured, and even if the bottleneck size is 1. While BREATH could be extended readily to allow for multiple samples per host, this is not implemented in the current version.

We use the terminology “ancestral to the sample” (ATTS) to represent hosts who have at least one sampled host amongst their descendants. (Sampled hosts are technically considered “ancestral” to themselves for the purpose of the nomenclature, even if they have no sampled descendants.) The sampled descendant host may not be next in the transmission chain, and can be separated from the ATTS host by any number of sampled or unsampled hosts. All hosts of all types in our trees are ATTS; we do not model “unknown unknown” hosts (unsampled hosts with no sampled descendants). “Multiply ancestral to the sample” (MATTS) refers to hosts who are direct sources of at least two hosts that are ATTS. The distinction between IUHs and members of chains is that the former are always MATTS and the latter never are. Sampled hosts may or may not be MATTS.

A MATTS host must colour at least one internal node of the phylogeny, because the pathogen common ancestor of the samples transmitted to both of its child hosts must have existed during its infection. The converse is also true for unsampled hosts: if they colour an internal node then they must be MATTS. Thus the number of IUHs is bounded above by the number of internal nodes of the phylogeny, which for *N* samples is equal to *N* − 1. The number of hosts in unsampled chains, on the other hand, is unbounded. Each chain consists of 1 or more unsampled, non-MATTS hosts and the chain ends when one of these infects a host that is either MATTS or sampled. (This distinction was previously alluded to by [23], in section 3.2 of the appendix.) In this way, we allow for an arbitrary number of unsampled individuals. Each sampled host and IUH is associated with a colour. We require that the section of the tree corresponding to each individual host colour (except the colour corresponding to chains) be continuous – in other words, colours are not broken up or interrupted by other colours. This corresponds to assumptions that hosts are not superinfected or reinfected and the bottleneck at transmission is complete.

### 2.2. Bayesian decomposition

#### Notation

Let *G* be the genealogy of the pathogen, a timed phylogenetic tree, and *T* be the transmission tree. The colouring of the nodes of *G* encodes *T* : it describes who infected whom and when, and where there are chains of unsampled transmission that are ancestral to the samples. *T* and *G* are the two main objects that we want to infer. We also have a number of parameters: *θ* are the parameters describing the underlying epidemiology. These include the mean number of secondary infections (if transmission is not interrupted by sampling or the end time) *C*^tr^, and the shape and rate parameters for the times to transmission and sampling (*A*^tr^, *B*^tr^ for transmission and *A*^*s*^, and *B*^*s*^ for sampling), and the sampling fraction *q*. Shape and rate parameters define the gamma distributions for times to infection and times to sampling. We also allow a parameter and corresponding prior for the time of origin of the outbreak (which is different from the root time of the phylogeny, because the index case could be infected before the first branching event). A list of model parameters is given in Table 1.

**Table 1:**
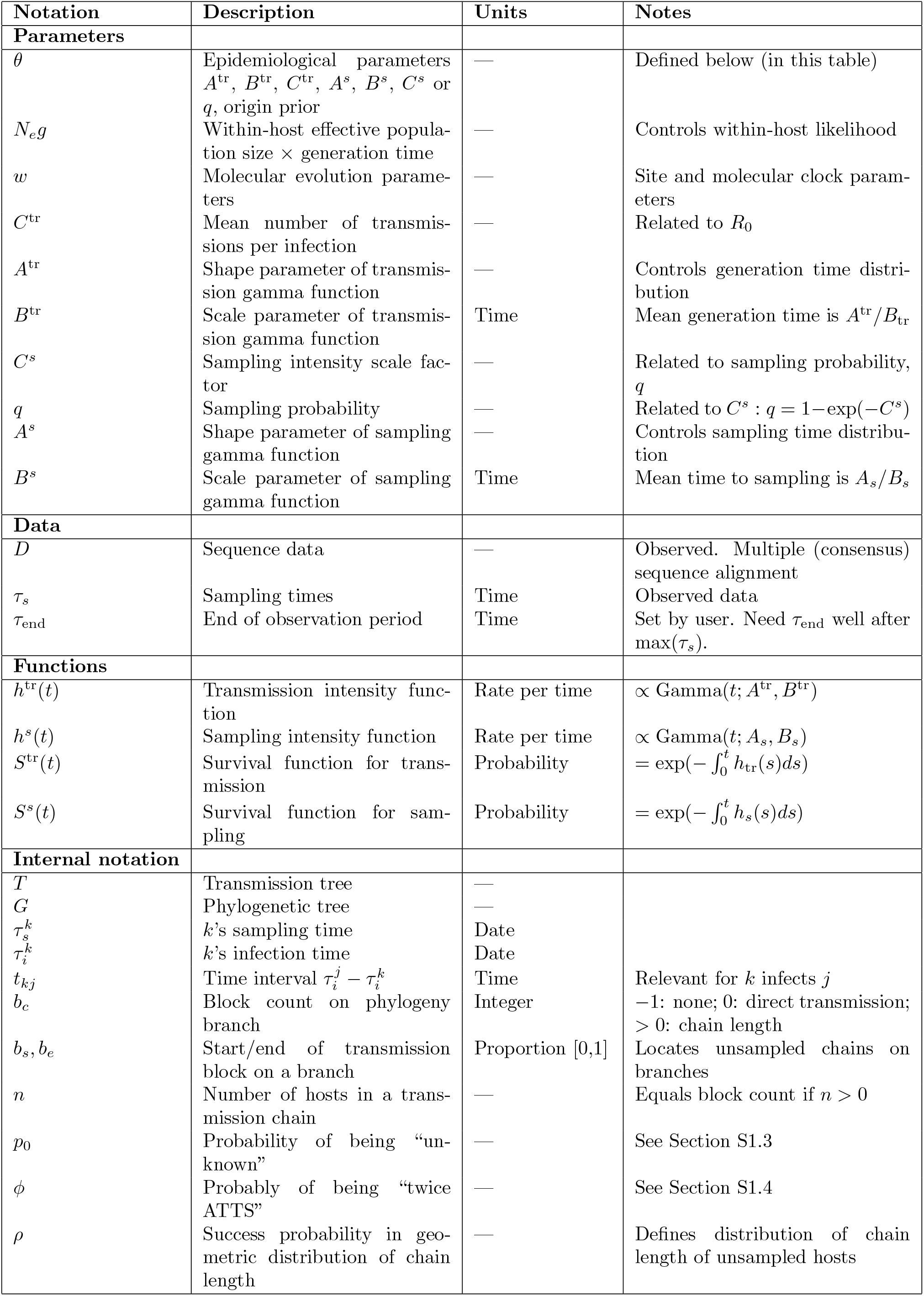
BREATH notation and parameters.

*N*_*e*_*g* is the within-host effective population size multiplied by the within-host generation time, and *w* represents the parameters describing the evolutionary model, including the molecular clock. (The within-host generation time *g* is that of the demographic process within each host; if *T* describes spread between organisms it will usually be that of the pathogen, whereas if *T* instead describes spread between locations such as farms, it may be the serial interval of transmission among individuals within a location.) We assume a strict bottleneck: only one lineage is transmitted at infection.

We have sequence data *D*, and sampling times *τ*_*s*_. The decomposition is essentially the same as it is in the BEASTLIER model [12], with posterior probability:

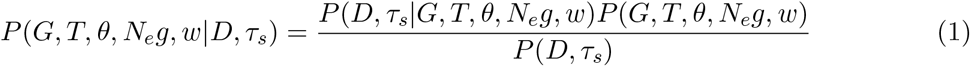

where as usual we do not know the denominator. Samples from this posterior can be drawn using MCMC.

We have, for the numerator,

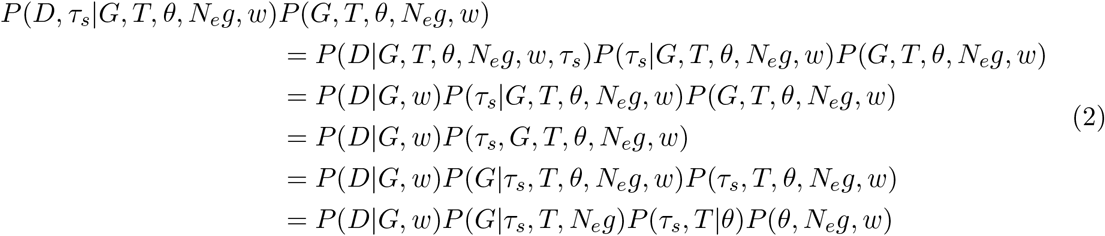

The third line of Eq. (2) follows from the assumption that conditional on *G* (the genealogy), the sequence data are independent of *T*, the parameter *N*_*e*_*g*, the sampling times and the epidemiological parameters. The last line assumes that the genealogy *G* does not depend on the molecular evolution model *w*, and the transmission process does not depend on the within-host parameter *N*_*e*_*g* (it is entirely epidemiological).

In the last line, *P* (*D*|*G, w*) is the likelihood of the sequence data given the phylogeny and *w*. This likelihood can be computed with Felsenstein’s pruning algorithm [1] in the usual way. The next term *P* (*G*|*τ*_*s*_, *T, N*_*e*_*g*), as in previous work [9], describes the likelihood of the genealogy given the transmission tree, within-host parameter and sampling times. This is a product of the likelihoods of smaller (independent) genealogies occurring in different hosts. Here we mainly take the same approach as in previous models [9, 17, 35, 28], and use a constant-size coalescent model to define the likelihood of a within-host tree, conditioning on the lineages coalescing after the time of infection (forward in time) such that only one lineage can be transmitted in a transmission event (see Supplementary Materials).

We need to compute *P* (*τ*_*s*_, *T*|*θ*), which is the heart of the BREATH model. The last term in the last line, *P* (*θ, N*_*e*_*g, w*), is the prior. Putting these ingredients together, we have

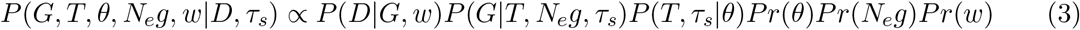

### 2.3. Epidemiological likelihood

We wish to find *P* (*T, τ*_*s*_|*θ*) in (3). This calculation is one of the novelties in BREATH, because it requires minimal changes to the model parameters despite potential wide variation in the number of unsampled cases, and allows us to account for the right truncation that occurs in a real-time outbreak. (The other novelty is the MCMC moves and BEAST implementation that enable simultaneous inference of the phylogeny and transmission tree using the colouring.) The parameters in *θ* represent parameters describing the transmission and sampling processes and the time at which observation ends, *τ*_end_. We do not infer *τ*_end_. *τ*_end_ must not be chosen to be equal or too close to the final sampling time, max(*τ*_*s*_), because this violates assumptions in the decomposition and leads to erroneous results; see Sections S6 and S8. If *τ*_end_ is not known, we suggest using the date when the analysis is being performed (unless analysis is initiated because of the arrival of a new case, in a real-time context).

The likelihood of the transmission tree and sampling times given the epidemiological parameters, *P* (*T, τ*_*s*_ |*θ*) in (3), can be written recursively. We use 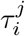 to denote *j*’s time of infection, and 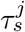 for *j*’s time of sampling. If *j* is not sampled, there is no sampling time; in this case we still compute a likelihood, but it describes the likelihood that *j* was not sampled given *j*’s infection time. Accordingly, set *s*_*j*_ be the sampling event (either sampling at time 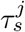, or unsampled) for host *j*. We let *T* (*k*) refer to the transmission subtree rooted at host *k*, and use *j →k* to mean “*j* infected *k*”. We start with the index (root) host *r*. We have

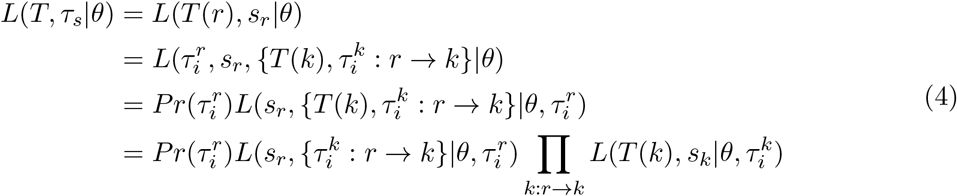

Here, 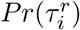 is the prior on the time of infection of the index host. We use an (improper) uniform prior by default. Improper priors are problematic if there is a need to sample from the prior, and when estimating marginal likelihoods through path or nested sampling, but are usually not a problem for sampling the posterior, since the age of the tree is driven by the rest of the prior and by the transmission and sampling intensity functions (see below and *Implementation*, Section 2.4).

The index host is the sampled host or IUH associated with the root of the phylogeny; we assume that if unsampled, it is MATTS. The terms 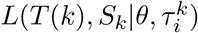 will in turn become a product of the likelihood of *k*^*′*^*s* events (sampling and infecting others, or not, and the product of the likelihoods 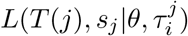 for the *j*s that *k* infects); infectees may start chains of unsampled transmission. We still need the terms 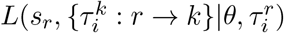 for individuals and for unsampled chains. To write these explicitly, we use a recurrent events model, and we treat individual hosts and chains of unsampled transmission separately.

#### Transmission and sampling

We use a recurrent events approach with two kinds of events: sampling, and transmission to another individual. Each has an intensity function, *h*^*s*^ and *h*^*tr*^ respectively. For each intensity function,

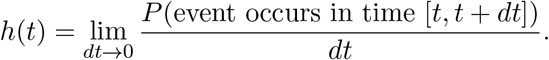

In a recurrent events model, the probability density for *k* events at times *t*_1_ *< t*_2_ *<* … *< t*_*k*_ with intensity *h*(*t*), in an interval from 0 to *t*, is

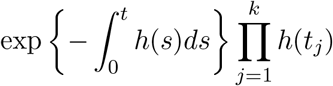

(see [3]). The intensity functions are like hazard functions, except that the event of interest can occur more than once. The expression in the exponential is the probability that the event does not happen in the interval [0, *t*]. It is analogous to a survival function. We will denote these as *S*, with

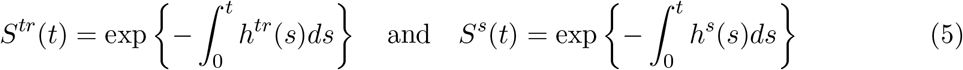

for transmission and sampling respectively. In this version we sample each host a maximum of once, but the model could readily be extended to multiple sampling events.

As an example, an individual infected at time 0, who infects another at time *τ*_1_ and is sampled at time *τ*_2_ would have a likelihood contribution of

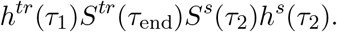

Since individuals must be infected before infecting others or being sampled, *h*^*tr*^(*t*) and *h*^*s*^(*t*) must be supported only for *t >* 0. We use the gamma shape to allow considerable flexibility in our assumptions about how fast these processes occur: *h*^*tr*^(*t*) = *C*^*tr*^gamma(*t, A*^*tr*^, *B*^*tr*^), and *h*^*s*^(*t*) = *C*^*s*^gamma(*t, A*^*s*^, *B*^*s*^). *C*^*tr*^ is the mean number of new infections that an infectious individual is expected to cause over the course of their infection if their infections are not interrupted by the end of the study or by sampling (similar to the basic reproductive number *R*_0_). *C*^*s*^ scales the transmission intensity function. The overall eventual sampling probability is 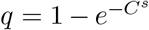 (1 minus the probability that sampling never occurs). *A* and *B* refer to the shape and scale parameters respectively. These functions are the building blocks of the likelihood, as they define the contributions from individuals infecting others (or not) and being sampled (or not).

#### Right truncation

BREATH accounts for the possibility that an outbreak might not be over at the time of analysis. This can create bias, because faster-occuring events are more likely to be observed. Here, we only have individuals in our tree if they are ATTS (recall that sampled individuals are ATTS to themselves): if *X* and all of *X*’s descendants are not sampled before the end time of our study (*τ*_end_), then we never know anything about *X* at all. This means that our data are right truncated. This is distinct from censoring (under censoring, we know about an individual, but the event of interest did not occur in our observation window). If the density for observing an event at time *t* since infection is *f* (*t*), but we know that we could only have observed this individual if *t < Y*_*R*_ where *Y*_*R*_ is a right truncation time, then the appropriate contribution to the likelihood for this individual is 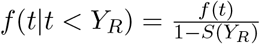 (see [2], Chapter 3).

We begin with the individual host (sampled or IUH). We need to know the appropriate *S*(*Y*_*R*_). Individuals are only in the model at all if they are either sampled or ATTS by the end time *τ*_end_. Let the intensity function for the event: “either infects someone ATTS or gets sampled by *t*” be *h*^*E*^(*t*), and let the survival function be *S*^*E*^.

In a time [*t, t* + *dt*), in our model at most one event can happen (*dt* is very small and rates are finite); either the individual can be sampled, or they can infect someone. If they infect someone, for our event (ATTS) to occur, that person has to eventually become ATTS.

Let *p*_0_ be the probability that an infectee and all of their descendant infections are unobserved – the probability of being an “unknown unknown”. We derive *p*_0_ with standard methods (see Supplementary Materials). Once we know *p*_0_, we have

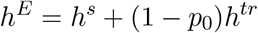

and therefore

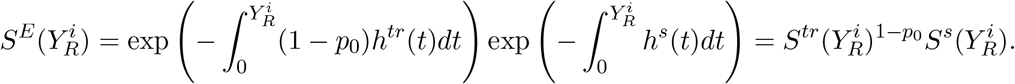

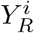, the right truncation time for host *i*, is the time between *i*’s infection 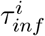 and the end of the observation period, 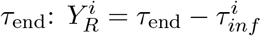.

#### Likelihood for individual hosts

We need to build the terms 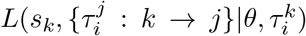 to have an explicit likelihood for the transmission tree in (3). For an individual host *k* who infects some others, indexed by *j*, we build this likelihood using the intensity *h*^*tr*^ at which *k* infects others, and the intensity *h*^*s*^ for *k* being sampled.

Throughout, *τ* values will be in “calendar time” and *t* will denote time intervals. Let *t*_*kj*_ be the time interval between *k*’s infection and 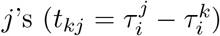, and let 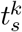 be the time between *k*’s infection and sampling. The contribution to the likelihood that comes from infecting others has a term 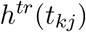 for each host *j* that *k* infects, along with a survival-like term, 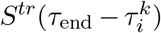 or 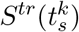, reflecting the fact that no other infection events happened during the time that *k* was a potential infector. If sampling prevents onward transmission (for example because hosts are treated effectively or isolated, and no longer infect others), then *k* stops being a potential infector of others when *k* is sampled. Otherwise, *k* could potentially infect others until *τ*_end_, though late infections are often very unlikely (as determined by the intensity *h*^*tr*^). Let the length of time when *k* could infect another be

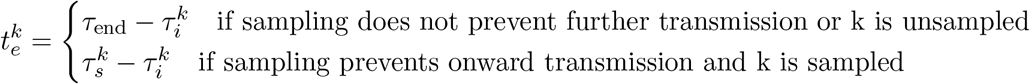

The transmission process gives a contribution

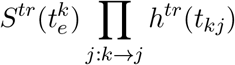

to 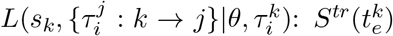 for no transmission events happening at times other than 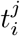 throughout the time when *k* could infect others, and for each transmission event that does occur, a term *h*^*tr*^ for its likelihood. If *k* does not infect any others, there is no product term.

Similarly, let the relevant time interval for *k*’s sampling be

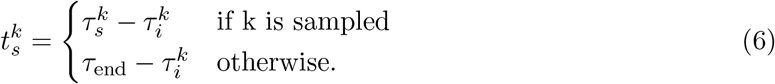

The sampling process has a term with 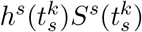 if *k* is sampled, and only 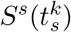 otherwise. Putting this all together, we have the likelihood for an individual host *k*’s time of sampling (if sampled) and of infecting others:

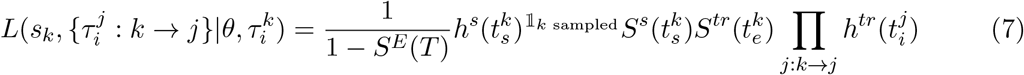

This is the main ingredient in (4). The term in the product in (4) breaks down recursively into a collection of terms like this one, for individual hosts, and terms for the chains of unsampled transmission, which we treat next.

#### Likelihood for unsampled chains of transmission

We now derive the likelihood for the chains of unsampled individuals. Such a chain is initiated when an infector infects someone who is not sampled and who is precisely once ATTS. Subsequent cases in the chain are unsampled, and the chain persists until one of two things occurs: either an infectee is sampled, or an infectee is MATTS (multiply ancestral to the sample; see above). That final infectee is not part of the chain even if they are unsampled, because in that case, their infection must contain at least one internal node of the phylogeny. The rightmost grey host in Figure 1 illustrates this: the chain of length 3 ends because this host is MATTS (infecting both B and another chain). Its infection contains a phylogenetic node. Also, the index case in Figure 1 hosts an internal phylogenetic node, has two lineages that are ultimately ATTS and so is MATTS (ATTS in two ways in this case: through *C* and the chain). This chain ends when *E* is infected, because *E* is sampled. A chain has three parameters: the number of hosts *n* (*n* = *b*_*c*_, the block count, if *b*_*c*_ *>* 0), the duration *t* of the chain, and the right truncation time. In the product term 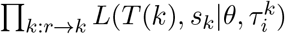 in (4), suppose *k* is a chain that ends when host *m* is infected. We have

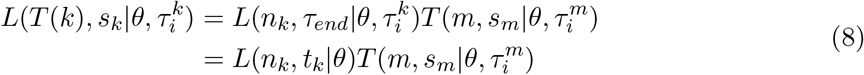

with chain duration 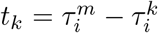 and number of hosts *n*_*k*_. We need *L*(*n*_*k*_, *t*_*k*_|*θ*). We drop the subscript *k* in what follows.

The number of unsampled infections in the chain is geometric, because for each infectee *X* in the chain, there is a probability that *X* either is sampled or *X* infects someone who is MATTS (ending the chain). The number of infections in the chain is the number of trials that happen before one is successful, which means *n ~* Geom(*ρ*). We need the success probability *ρ*. We will use the same idea we used for *h*^*E*^ above to find *ρ* and also to manage the right truncation that occurs because we cannot observe a chain at all unless a MATTS individual is infected by the chain before the observation period ends. We proceed by finding *ϕ*, the probability that a host will eventually be MATTS, and then *ρ*. We condition on the block forming at all by dividing by *P*_1_, the probability that the initial infecting lineage is exactly once-ATTS. The block likelihood is therefore:

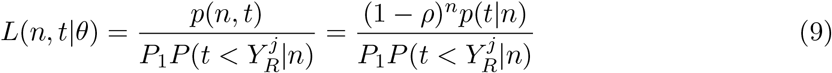

where 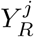 is the chain’s right truncation time and *t* is the duration of the chain. The numerator contains the geometric probability for the *n* cases in the block that were not MATTS. The *n* + 1’st is MATTS, but their sampling and transmission information is treated not with *ρ* but with the individual likelihood. The density for the block duration given *n, p*(*t*|*n*), is the density for the sum of *n* transmission times: *p*(*t*|*n*) *~* gamma(*nA*^*tr*^, *B*^*tr*^). The term 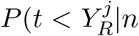 is the CDF of this gamma density. See the Supplementary Materials for the computation of *ρ* and *P*_1_.

Figure 2 shows a schematic of a small outbreak, and connects the key mathematical expressions with the events in this outbreak. Table 1 lists the notation and parameters.

**Figure 2:**
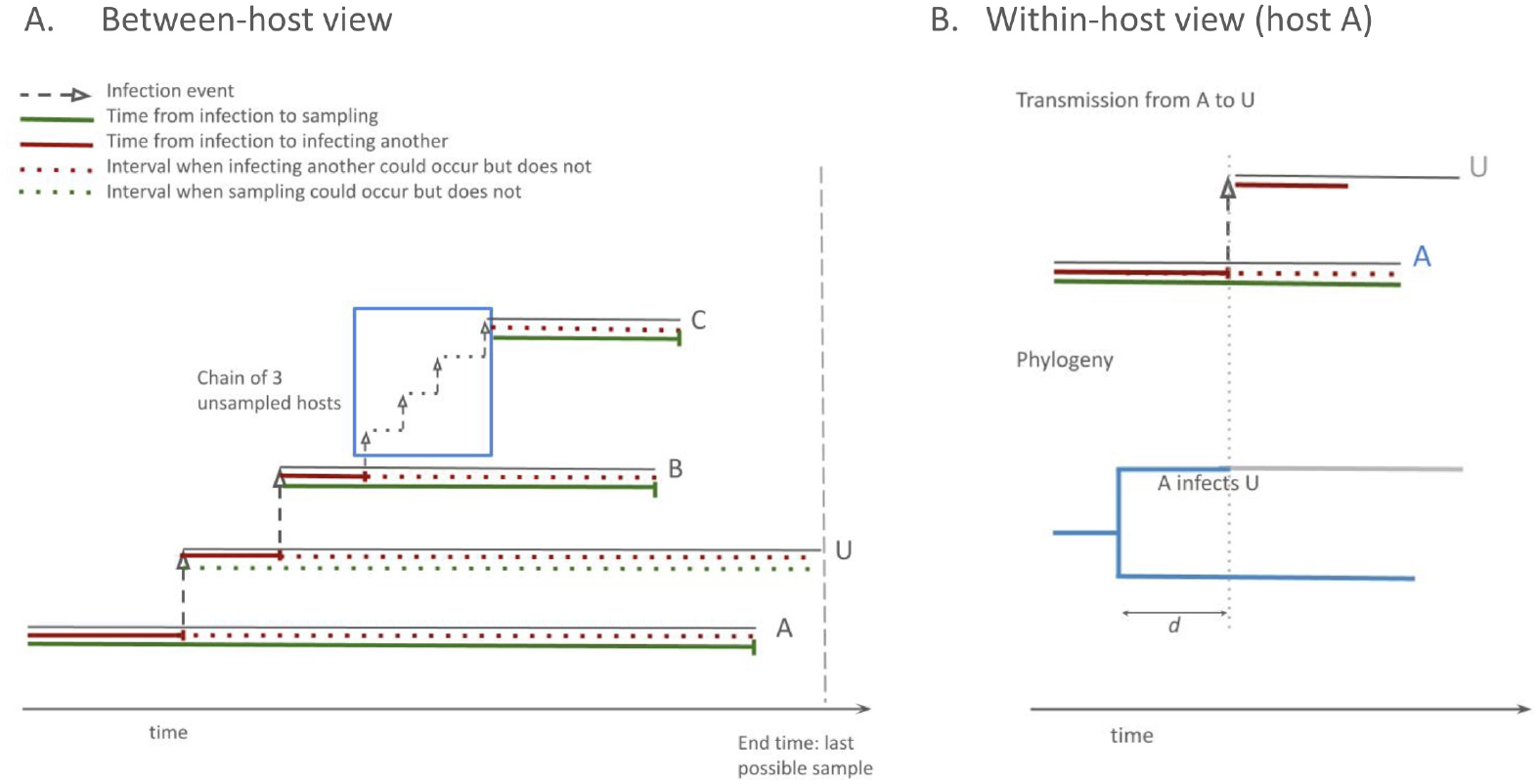
A small outbreak. A (left): Hosts A, B and C are sampled (vertical green markers). A infects U (unsampled), who infects B. Before being sampled, B infects a chain of three unsampled hosts. This chain infects C, who is sampled. Solid lines show the times between infection and sampling (green) and infection to infecting another (red). A solid red line lasting a duration *t* corresponds to a term *h*^*tr*^(*t*)*S*^*tr*^(*t*) in the likelihood, because no transmission event happens during the solid period *and then one does at the end*. A dashed red line lasting duration *d* corresponds to a term *S*^*tr*^(*d*) in the likelihood, and there is no *h* term because no event happens. A solid green line with duration *l* corresponds to a term *h*^*s*^(*l*)*S*^*s*^(*l*) and similarly a dashed green line of duration *d* corresponds to a term *S*^*s*^(*d*), with no *h* term. For each process we condition on the fact that there is a last possible time when anyone could have been sampled (vertical dashed line). This means, for example, that case C must have been observed before the end time, so any terms in the model for C are conditioned on a right truncation time which is the time between when C is infected and the last possible sampling time. The same conditioning is done for other cases, but it makes less of a difference for A who was infected earlier than C. B: (right). Within-host view focusing on the phylogeny within host A (blue). When A infects U, the lineage that is transmitted changes colour. A host harbours two lineages at the same time for a duration *d*. The intuition for how *d* affects the likelihood is described in Equation (S1.1). When *N*_*e*_*g* is small, shorter durations are much more likely than when *N*_*e*_*g* is larger.

### 2.4. Implementation

The transmission tree likelihood, the MCMC proposals, a simulator and some post-processing tools are implemented in the `BREATH` package for BEAST 2 [22].

The BEAST 2 implementation associates between one and three parameters with each branch *b* of the phylogeny. There is always a “block count” *b*_*c*_, for which there are three modes of interpretation (see Figure 1). Each mode dictates the existence and distinctness of a block start *b*_*s*_ and a block end *b*_*e*_. The *b*_*s*_ and *b*_*e*_ define positions between start and end of *b*, so, if *b* has length *b*_*k*_, 0 *≤ b*_*s*_ *≤ b*_*e*_ *≤ l*(*b*):

- *b*_*c*_ = *−*1 represents no infection happening on this branch. Neither *b*_*s*_ nor *b*_*e*_ exist. The host associated with the phylogenetic nodes at both ends of the branch must be the same. Conversely, if the hosts at both ends of a branch are the same, by the continuity (no reinfection) requirement we must have no unsampled hosts on the branch, and *b*_*c*_ = *−*1. If there were reinfection, such that for some hosts, multiple distinct sequences were available, the separate infections could be included as distinct taxa and labelled as if they were distinct hosts.
- *b*_*c*_ = 0 represents that a single infection took place on the branch, and the host at the top of the branch infects the host at the bottom of the branch, so there are no unsampled hosts in between. *b*_*s*_ nor *b*_*e*_ exist, but are equal.
- *b*_*c*_ *>* 0 represents the presence of a chain of unsampled hosts, and as the duration of this chain is nonzero, we must have *b*_*s*_ *< b*_*e*_.

We initialise *b*_*c*_ to 0 and *b*_*s*_ = *b*_*e*_ = 0.5 for each branch, forming a valid starting state.

### 2.5. Dimensionality

We associate each point in the phylogeny either with a sampled host, an IUH, or a chain of unsampled transmissions. We associate each sampled host or IUH with a colour. (There are a maximum of 2*N* colours for a tree with *N* taxa: *N* sampled hosts, up to *N* − 1 IUHs and a colour for chains.) Each edge of the phylogeny is given a number *b*_*c*_ encoding the number of transmissions on the branch. Each edge has up to 2 other parameters (*b*_*s*_ and *b*_*e*_ above). This constrains the dimension of the phylogenic tree object as the number of unsampled hosts varies. The minimum number of block starts and ends is *N* −1, when there are no unsampled hosts and *N* infection events. All, except the infection of the index, occur on a branch with *b*_*c*_ = 0 and all other branches have *b*_*c*_ = *−*1. The maximum is 4*N* − 4, where every sampled host’s infection event occurs on the terminal branch leading to their tip, no two internal nodes have the same colour, and every block has nonzero length. An additional parameter is always required for the root edge (this allows that the index host, who is always either sampled or an IUH, does not necessarily have a coalescent event at the moment of their infection). Simple reversible-jump moves are required to change the number of block-related parameters on a branch between 0 and 2 as *b*_*c*_ changes (see the Supplementary Materials), but we avoid the need for the much more computationally challenging reversible-jump approach of TransPhylo [17], where every unsamped individual is a distinct host with a time of infection and time(s) of infecting others, and where the total number of these is unbounded.

### 2.6. MCMC proposals

There are three MCMC operators that handle proposals for *b*_*c*_, *b*_*s*_ and *b*_*e*_: the infection mover, the block boundary operator, and the block addition and removal operator. Both of these can change the various values of *b*_*c*_ such that the dimension of the parameter space also changes, and as such may be reversible-jump moves. In what follows a “dimension change” is a situation where a *b*_*c*_ changes from −1 to 0, −1 to 1 or more, 0 to 1 or more, or (in each case) vice versa. A redraw of block boundaries for a branch *b* with *b*_*c*_ = 0 consists of drawing a single value *v* uniformly from (0, 1) and setting *b*_*s*_ = *b*_*e*_ = *vl*(*b*). If *b*_*c*_ *>* 1 then two values *v*_1_ and *v*_2_, such that *v*_1_ *< v*_2_, are drawn from the unit triangle and then *b*_*s*_ set to *v*_1_*l*(*b*) and *b*_*e*_ to *v*_2_*l*(*b*). For details of the Hastings ratios, see the Supplementary Materials.

The infection mover, as the name suggests, picks an infection and moves it elsewhere. There are two versions.

- The “narrow” version picks a branch *b* with *b*_*c*_ *≥* 0 and moves an infection from its block to the block of its sibling branch. If this leads to a violation of colouring rules then the move fails. If a dimension change on each branch has occurred, then block boundaries are redrawn based on the new *b*_*c*_ values. The move is reversed by choosing the sibling as the initial branch.
- The “wide” version does the same except that the destination branch is not required to be the sibling of *b*. Instead it is chosen from the complete set of branches in the tree with probability proportional to its length.

The block boundary operator selects a branch *b* uniformly (without taking branch lengths into account). If *b*_*c*_ = *−*1 the move returns without doing anything else. Otherwise block boundaries are redrawn on *b*.

For the block addition and removal operator, removal is chosen with 50% of the probability. Not all infections can be removed without leaving the transmissions in an invalid state, for example, a single infection on a branch between two sampled hosts cannot be removed. First, the set *B*_elig_ of branches for which an infection can be removed without causing such a problem is identified, along with the number of distinct block boundaries on them. If *B*_elig_ is empty then the move fails. Otherwise, an eligible branch is randomly selected. An infection in the block on the branch is removed as follows: if *b*_*c*_ = 0, reduce *b*_*c*_ to *−*1. If *b*_*c*_ = 1, reduce *b*_*c*_ to 0 and redraw a single block boundary. If *b*_*c*_ *>* 1, reduce *b*_*c*_ by one and leave the block boundaries unchanged. If we are instead adding an infection, let *B* be the complete set of branches of *T*. Select any branch *b ∈ B* with probability proportional to its length *l*(*b*). If *b*_*c*_ = *−*1, set *b*_*c*_ to 0, and if *b*_*c*_ = 0, set *b*_*c*_ to 1; then redraw block boundaries. If *b*_*c*_ *>* 0, increase *b*_*c*_ by 1 and leave the block boundaries unchanged. Changes to the phylogeny use the standard BEAST 2 set of operators, with one modification due to the variable dimension. A change in the length of a phylogenetic branch *b* is, if *b*_*c*_ *>* −1, actually a change to two or three different parameters, because, if *l*(*b*) becomes 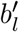 and *b*_*s*_ = *vl*(*b*), then *b*_*s*_ becomes 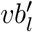 (and similarly for *b*_*e*_ if it is distinct). To deal with this, each of the standard operators has a wrapper that updates the Hastings ratio with an extra contribution before and after the tree proposal is performed: for each branch *b*, empty blocks (*b*_*c*_ = *−*1) do not contribute anything, blocks with *b*_*c*_ = 0 contribute *l*(*b*) and with *b*_*c*_ *>* 0 contribute *l*(*b*)^2^. The HR of the tree operator is multiplied by the product of individual branch contributions after the operation and divided by the total contribution before the operation.^1^ Furthermore a new state will be rejected if the colouring is invalid. There are some particular subtleties to why dedicated tree moves are not necessary that are worth expanding upon.

Firstly, in BEAST data structures a node is one of 2*N* − 1 distinct objects (for *N* sequences), annotated with its parent node and the length of the branch leading to it (if it is not the root). A prune and regraft move is accomplished by changing parents and branch lengths, not by cutting an existing branch into pieces and merging two others. This means that we can further annotate each node with the value of *b*_*c*_ for their branch, and these need not change when parents or lengths do. The root node in BEAST is not a fixed data object; instead “being the root” is a property that can be given to any internal node. Thus we can allow every node to have block parameters but simply ignore them while that node is the root.

Secondly, we do not need to annotate internal nodes with colours, as all such colours are implied by the tip colours and the blocks. Nodes separated by a branch *b* with *b*_*c*_ *>* −1 must have different colours. If a node is connected to a tip by branches all of which have *b*_*c*_ = *−*1 then it must have the same colour as that tip. We assume that the IUHs are interchangeable, so we do not need to differentiate internal nodes coloured by one unsampled host or another. This is enough to colour the tree without actually making colour into a node attribute, which means that specialised colouring-preserving moves are not necessary (unlike in BEASTLIER). There are some configurations which lead to colourings which are incoherent, notably those where two *tips* are connected by branches with only *b*_*c*_ = *−*1, but a proposal for such a state will simply be rejected.

Because the MATTS host associated with the root node is not assumed to have been infected at the phylogenetic TMRCA, we do require a root branch length defining the infection time of the index case, in common with BEASTLIER and also birth-death phylodynamic models [5]. This is a simple positive real number, proposals for which are standard.

For the population size parameter *N*_*e*_*g*, we used a standard Bactrian scale operator.

### 2.7. Parameters for the outbreak analysis

We analyzed a previously-described TB outbreak [7] with 86 detected cases in Germany between 1997 and 2010. Single nucleotide polymorphisms are available in the previous publication along with month and year. We used *A*^*s*^ = 10.0, *B*^*s*^ = 6.5 and *q* = 0.75 for the sampling probability. If an individual is sampled, the mean time to sampling would be *A*^*s*^*/B*^*s*^ = 1.5 years since infection, with a standard deviation of 0.5 years. For transmission, we used *A*^*tr*^ = 10, *B*^*tr*^ = 8.5 and *C*^*tr*^ = 2. This reflects the assumption that the mean number of transmissions if they were not interrupted would be 2, the mean time to sampling would be 10*/*8.5 = 1.17 years) with a standard deviation of 0.37 years. Note that these times are not adjusted for the end of the study period (that occurs in the right truncation adjustment, not in these parameters).

These values were informed by the analysis of the same outbreak with TransPhylo [17]. We used the bModelTest site model [16] and a strict molecular clock. We used a uniform prior [0, 10] for *N*_*e*_*g* (see Supplementary Materials). Other parameters are as specified in the XML file at https://github.com/rbouckaert/BREATH/releases/tag/v0.0.5.

### 2.8. Availability

BREATH is available at https://github.com/rbouckaert/BREATH/ with a tutorial at https://github.com/rbouckaert/BREATH/tree/main/doc/tutorial. The data for the TB outbreak are available in [7] with a demonstrative dataset of 40 sequences in the tutorial. The augmented phylogenies can be viewed using the interactive who infected whom package https://github.com/Lars-B/interactive-wiw or viewed in `icytree`. To extract transmission trees with who infected whom and when information, including individual unsampled hosts and blocks, there is a `breath-helper` tool in the `pyccd` package by Lars Berling at https://github.com/Lars-B/pyccd.

### 2.9. Applicability

BREATH is a method for the reconstruction of person-to-person transmission events in well-sampled outbreaks, using pathogen sequences as a key input (along with knowledge of the likely times between infection and sampling, and infection and infecting others, though these may be variable and imperfectly known). BREATH is, accordingly, not a suitable analytical tool for genomic surveillance data with very low sampling fractions. For example, consider a viral outbreak in which all known cases were tested, but only a random, uniform fraction *p* of individuals had their samples sequenced and included in the relevant dataset. For a transmission pair in which *A* infects *B*, both *A*’s and *B*’s sequence are in the dataset with probability *p*^2^. If *p* is small, there are likely to be too few direct transmission events for BREATH or any other person-to-person transmission reconstruction to be useful. Similarly, if only a fraction *q* of known cases in the outbreak were tested and all tests correspond to sequences in the dataset, *A* and *B* are both present with probability *q*^2^. It may be the case that despite overall low sampling in a jurisdiction, there are densely-sampled transmission clusters in the dataset. In this case these may usefully be analyzed with BREATH.

BREATH’s applicability also requires that sequences be informative of transmission. For example, if too many sequences are identical, phylogenetic methods of any kind will not be helpful. It is very unlikely that the opposite extreme will be relevant: at the person-to-person time scale, it is unlikely that there would be too much between-host genetic variation for phylogenetics to be a useful tool, nor (with pathogens like viruses or bacteria) that there is too much reassortment, recombination, or horizontal transfer of genetic material for phylogenies to be relevant. However, if a phylogenetic tree is known not to be a good model for the joint evolution of the sampled sequences, then BREATH and any other phylogenetic reconstruction will not be appropriate. Figure 3 outlines when BREATH is likely to be an appropriate tool. This will not be for the majority of genomic epidemiology studies, because densely sampled and sequenced outbreaks are the exception rather than the rule. In short, BREATH reconstructs person-to-person transmission histories and requires (1) sufficiently high sampling *of the transmitting individuals in a transmission cluster*, (2) that pathogen genomes be at least some what informative of transmission, and (3) some knowledge of the relevant time distributions (knowledge of individuals’ times of infection is not required). See Section S8 for more details.

**Figure 3:**
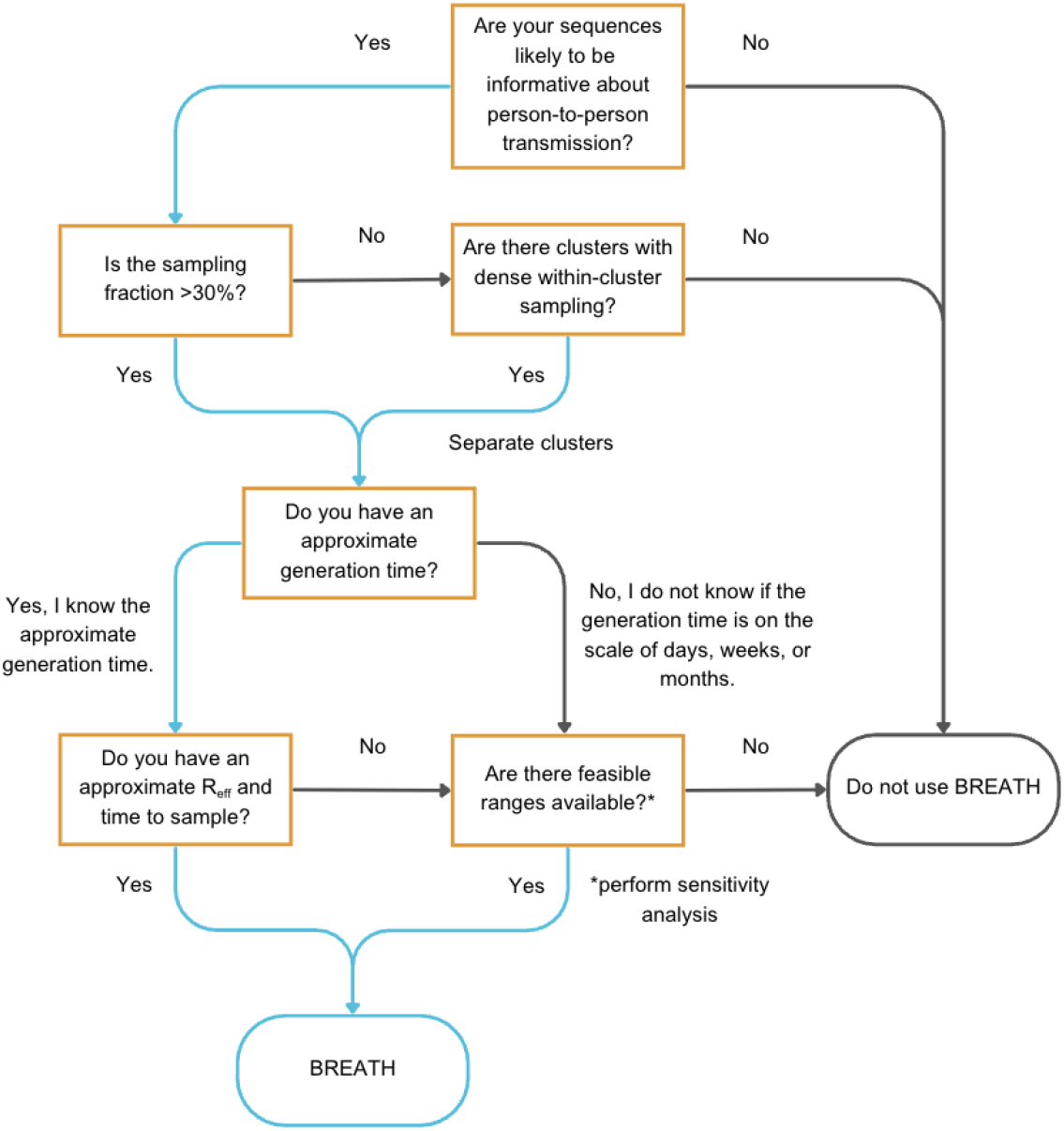
A decision tree to help determine whether BREATH is an appropriate analytical approach. If sequences are not likely to be informative of transmission, or if there are not clusters with reasonably dense within-cluster sampling, person-to-person transmission reconstructions from pathogen genomes are not feasible with any tool, including BREATH.

## 3. Results

### 3.1. Validation

We performed a well calibrated simulation study [36] to provide some confidence that the implementation is correct. To this end, we first sampled 100 transmission trees with 32 taxa using the `TransmissionTreeSimulator` app (details in Section S3). For the sampling intensity, we used a shape *A*^*s*^ of 10, a rate *B*^*s*^ of 5 and a sampling probability *q* of 0.75. For the transmission intensity, we used a shape *A*^*tr*^ of 10, rate *B*^*tr*^ of 8 and expected infection count *C*^*tr*^ of 2. The *N*_*e*_*g* parameter size was set at 0.5.

Further, we simulated an alignment on the tree with 2000 sites under an HKY model with *κ* = 2 and equal frequencies (no gamma rate heterogeneity) and a clock rate of 0.25. Then, we ran analyses for each of the 100 transmission trees in BEAST2 where the tree, block count, block start and end as well as the population size were estimated with BREATH. We measure how often a true parameter value is in the 95% highest probability density (HPD) interval, which for 100 runs should be in the range 91 to 99 to be acceptable. The coverage of tree length is 96, tree height 94, and infection count 95, so all in the acceptable range.^2^

We want to verify that if the model predicts that host *A* is infected by host *B* with probability *p* that the true probability under the model is indeed *p*. Figure 4 shows the who-infected-whom predicted probabilities and their “true” probabilities (how often events occurred in the simulated ground truth). Note that the size of each bin, corresponding to the number of predictions with the given probability of being infected by a particular host, decreases with increasing probabilities. So, there are many very low confidence predictions, and the higher the confidence, the lower the number of predictions. Ideally, all bars should cross the black x-y line, indicating that predicted probabilities equal actual probabilities. For the simulation study using the `TransmissionTreeSimulator` app, most bars are very close to the ideal, though the bar for 70-80% is perhaps just under what they should be, which may be due to the inherent stochasticity of the process. Note that a large number of the predictions in the 70-80% category are for being infected by an unsampled host, which are not usually the predictions of interest.

**Figure 4:**
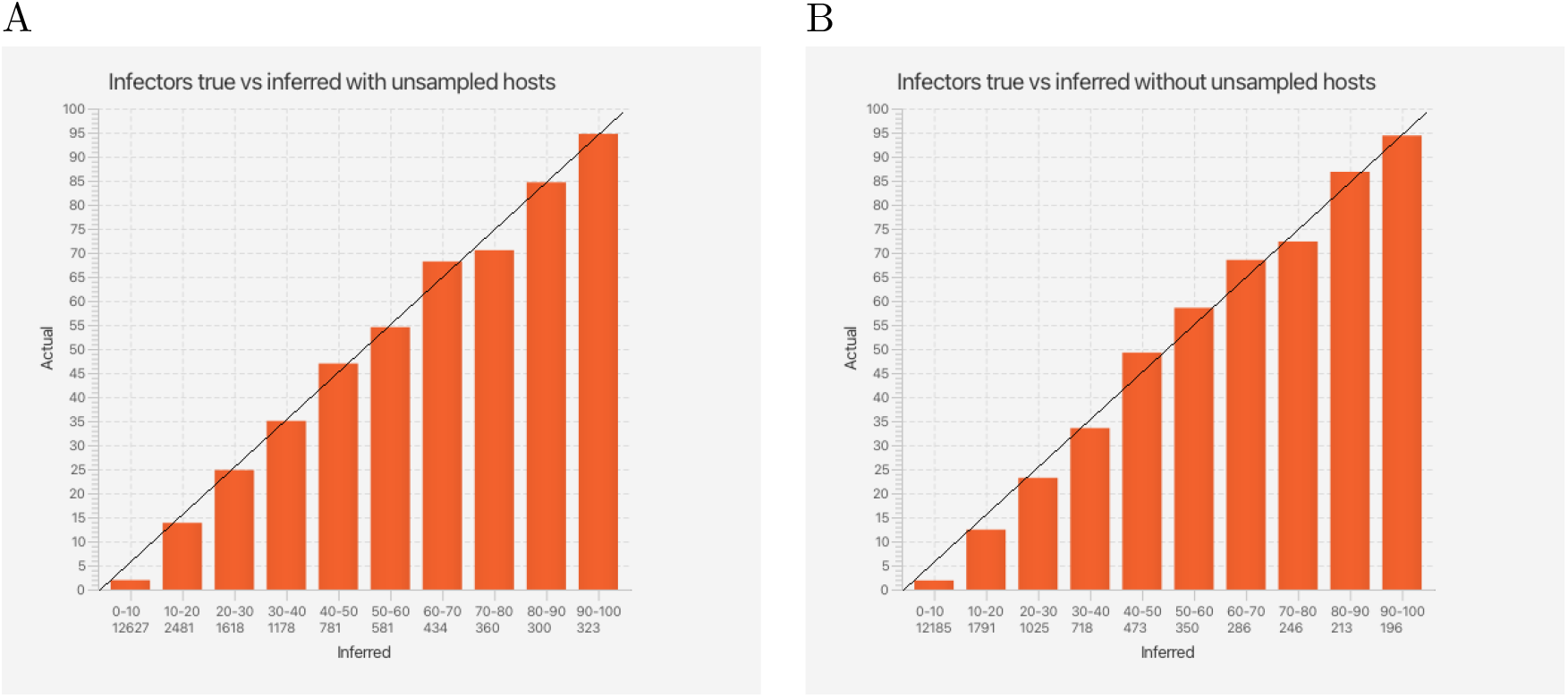
Who-infected-whom inferred probabilities versus true values for A with and B without unsampled hosts for simulation study with trees simulated using the `TransmissionTreeSimulator` app. The graph is constructed by counting how many inferred probabilities fit in a bin, then calculating how many of the true values are in a certain bin. Each bar represents a 10% sized interval, where the first bar represents predictions in the range 0 to 10%, the second bar ranges from 10% to 20%, etc. The number below the range represents the number of predictions that fit in that range for each of the 100 posterior distributions. For example, there are 323 predictions in the range 90-100% (last column panel A). The size of the bar is the percentage of times the true prediction is in the bin with the associated prediction. All predictions are as expected.

Section S5 shows similar results under a different set of parameters more suitable for the omicron variant of SARS-CoV-2.

#### 3.1.1. Sensitivity analysis

To explore how sensitive the analysis is to parameters of the hazard functions, we reran the analyses with misspecified parameters. Where the well calibrated simulation study uses *A*^*tr*^ = *A*^*s*^ = 10, *B*^*tr*^ = 8.0, *B*^*s*^ = 5.0, *C*^*tr*^ = 2.0, and *q* = 0.75, we reanalysed the data with deviations from these values as shown in Table 2. Parameter values were chosen to be compatible with the recommendations we outlined in Section S3.2. Increasing *B*^*tr*^ to over 8.5 led to zero posteriors at initialisation, so it was left at 8.0. A total of twelve variants were considered, for which 100 analyses were run, making a total of 2400 analyses. This is the second form of model misspecification that we check. The first was that the simulator was conditioned on producing 32 taxa, whereas the transmission likelihood cannot be so conditioned.

**Table 2:** Coverage of tree height, length and infection count under parameter misspecification in the simulation study. Simulations were done under the default parameters (*A*^*tr*^ = *A*^*s*^ = 10, *B*^*tr*^ = 8.0, *B*^*s*^ = 5.0, *C*^*tr*^ = 2.0, and *q* = 0.75). Values for the row “Original” are from inference under these (true) parameters. Values in the rows below are from inferences in which BREATH was given misspecified parameters (as listed). The infection count coverage is the most sensitive parameter to misspecification, and results depend most strongly on the sampling rate parameter *B*^*s*^ and transmission rate parameter *B*^*tr*^. The analysis is less sensitive to misspecification of the sampling fraction *q*, transmission shape *A*^*tr*^ and constant *C*^*tr*^.

|  | Tree<br>height | Tree<br>length | Infection<br>count |
| --- | --- | --- | --- |
| Original | 96 | 94 | 95 |
| $A^s = 9.0$ | 97 | 93 | **81** |
| $A^s = 11.0$ | 96 | 97 | **75** |
| $B^s = 4.0$ | 97 | 96 | **45** |
| $B^s = 6.0$ | 95 | 93 | **57** |
| $q = 0.55$ | 96 | 96 | 93 |
| $q = 0.95$ | 95 | **87** | **54** |
| $A^{tr} = 9.0$ | 96 | 94 | 96 |
| $A^{tr} = 11.0$ | 98 | 94 | 95 |
| $B^{tr} = 7.0$ | 95 | 95 | 92 |
| $B^{tr} = 8.5$ | 95 | 94 | 96 |
| $C^{tr} = 1.0$ | 96 | 97 | 96 |
| $C^{tr} = 3.0$ | 97 | 93 | 91 |

We calculated how well the tree height, tree length, origin and infection count were covered by the 95% HPDs. For the simulator based sensitivity analysis the origin was not available. Table 2 shows coverage of the parameters of interest, suggesting coverage is most sensitive to sample hazard parameters than transmission hazard parameters. In particular, coverage appears particularly sensitive to the sampling rate (*B*^*s*^) and less so sample shape (*A*^*s*^), and less sensitive to sampling constant (*C*^*s*^), transmission constant (*C*^*tr*^) and transmission shape (*A*^*tr*^). Table S1 summarises the who-infected-whom results in a manner similar to that shown in Figure 4, but in table format. Except for the rate parameters, most inferred who-infected-whom probability estimates are in the range X% to X+10% when the true cases are within X% to X+10%, but deviations exists, especially in the middle range. Who-infected-whom probability estimates are close to what is expected overall, but appear most sensitive to changes in sample rate parameter (*B*^*s*^), then the shape parameters (*A*^*s*^ and *A*^*tr*^), and less to the constants (*C*^*s*^ and *C*^*tr*^) and transmission rate parameter (*B*^*tr*^). Overall, estimates appear stable under misspecified hazard parameters.

### 3.2. Application to a tuberculosis outbreak

We analysed an alignment of 86 samples from a tuberculosis (TB) outbreak in Hamburg, Germany, described in [7]. We fixed the intensity function parameters as in the simulation study, and estimated the within-host coalescent parameter. Since we are not interested in estimating the site model (better done for tuberculosis with larger datasets comprising more evolution), we average over reversible models using bModelTest [16]. A strict clock is used for the branch rate model. To see what the impact is of the transmission tree likelihood on the phylogenetic tree, we compared BREATH’s results with BICEPS [27], a flexible coalescent prior taking epochs into account.^3^

Site models are indistinguishable across the two scenarios, which is as expected: the tree prior should not impact the way sequences evolve. However, the root height and tree length are somewhat lower in BREATH than in BICEPS: the mean height is 21.8 years for BICEPS with 16.2-28.9 years (95% HPD), and mean 15.4 years for BREATH (with 95% HPD 14.6-16.3 years). These correspond to a time of origin of early 1989 (late 1981 – mid 1994) in BICEPS, compared to mid 1995 (late 1994 – mid 1996) in BREATH. The published description of this outbreak listed a time of origin of 1993-1997 for what they termed the “Hamburg clone” (the main driver of this outbreak) [7].

The mean total tree length (sum of all branch lengths) is 217.6 years in the BICEPS model (146.4-304.3 95% HPD) and is lower at 169.2 years for BREATH (157.0-180.8 years 95% HPD). In both the branch length and the tree heights, we find a substantial reduction not only in the value but also in the uncertainty of the estimate. The change in node height estimates is most pronounced near the root, and not so pronounced at internal nodes that are nearer to the tips. Clade support differs between the two analyses (Figure 5A), also showing that the tree prior has a substantial impact on the inferred trees. Figure 5B shows an multi-dimensional scaling (MDS) plot, illustrating that while there are consistent differences between BREATH and BICEPS phylogenetic trees, these are comparable to the tree variability within each set.

**Figure 5:**
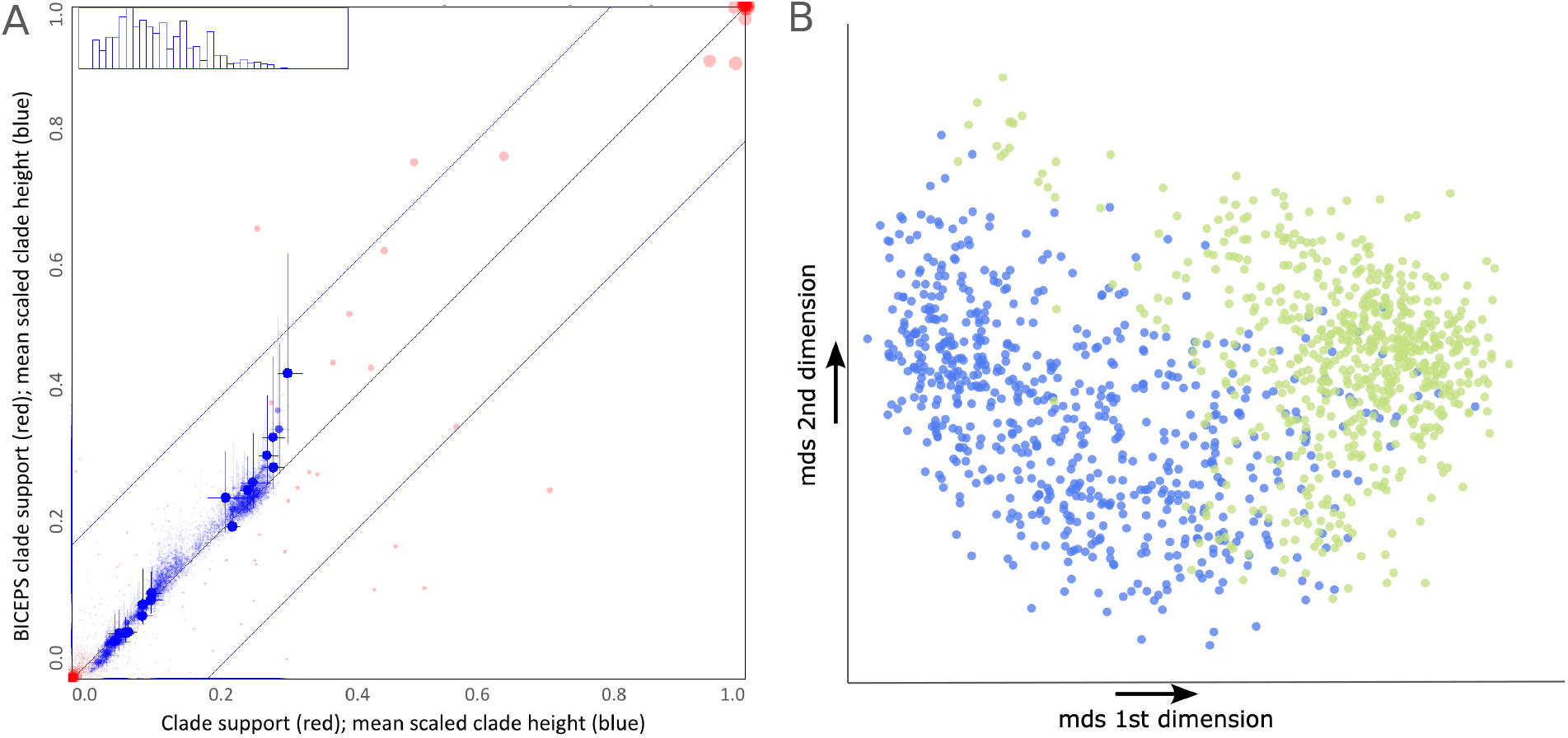
Difference between BICEPS and BREATH. A. Clade support (red dots) running from zero probability (bottom left corner) to 1 (top right corner) and scaled clade heights (blue dots). Dots are placed at the mean of the height of a clade. Both x and y axes are scaled so that the largest tree height (from among both data sets) is at the top right corner. Blue crosses indicate 95% highest probability density (HPD) intervals of a clade’s height. Larger dots have larger clade supports. We compare BREATH (horizontal axis) to BICEPS (vertical axis). The diagonal blue lines mark a 20% difference between BREATH and BICEPS: points within these lines have less than 20% difference, points outside have more, and can be considered as evidence of a substantial difference between the two posteriors. B. Multidimensional scaling plot generated by the TreeDist package in R [31] showing marked differences between the tree posteriors of the BICEPS (blue points) and transmission tree (green points) analysis.

Figure 6 shows BREATH’s inferred graph of who infected whom. Nodes in the graph represent hosts, and edges in the graph represent possible transmissions between hosts. The edges are annotated with probability of infection as inferred under the model. Hosts in tightly-connected parts of this graph have a high probability that they were infected by another sampled host (as opposed to an unsampled host); this probability is shown in parenthesis in Figure 6. These “clusters” are in areas of the phylogenetic tree where branch lengths are short, and sequences have short distances to each other; it is these short branches that likely drive the inference of infection by another sampled host.

**Figure 6:**
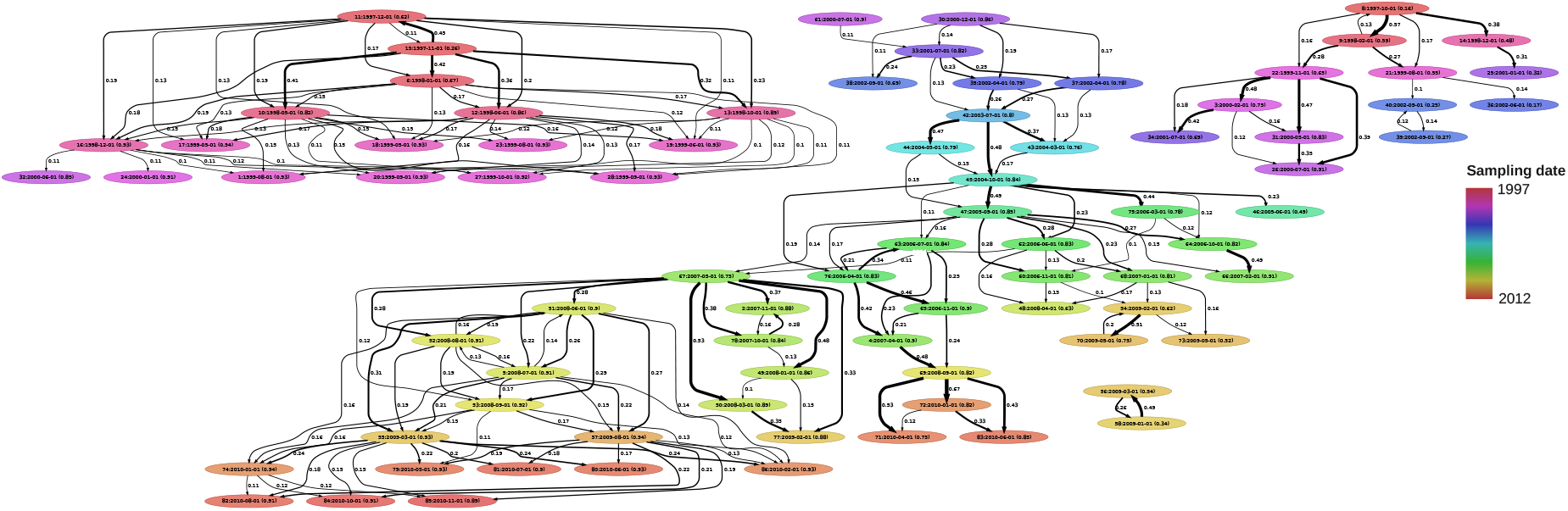
Who infected whom, predicted in TB outbreak. Numbers on branches represent probabilities the host at the tail of the arrow infecting the host at the head. Only the most important predictions (over 10% probability) are shown. Numbers in brackets next to labels are the inferred probabilities a host is infected by another sampled host. Hosts are coloured by time of sampling.

In our results, where pairs can be identified, the direction of transmission can be uncertain. Nonetheless, we find many high-probability direct transmission events, some individuals for whom the infector is not known and some individuals for whom there is a high probability that the infector was unsampled. There are three major sections of the transmission network. In the middle of Figure 6, there is a large transmission cluster spanning nearly the time frame of the data (2000-2010); in approximately 2003 there were only a very few individuals connecting the first and second stages of this component (hosts 42 – 45; see [7] Table S1, and then hosts 75 and 47 just after that; our cases are numbered in the order they appear in that table, i.e. case 1 is labelled “7199/99” in [7] Table S1). The top right and left components of the network are near the start of the outbreak (1997–2000), and corresponds to the lower two groups in the densiTree phylogenies in Figure 7. Despite the phylogenetic uncertainty, there are a number of relatively high-probability transmission events. Roetzer et al. [7] reported that while there were strong efforts to stop the outbreak, including closing a particular bar in 2006, the outbreak continued in Hamburg and spread to Schleswig-Holstein in 2006. Overall, we found an average of 17 unsampled cases with [10-23] 95% HPD. The average maximum block count was 2.16 hosts. This is plausible: according to the original paper, the outbreak spread primarily in a bar setting and impacted high-risk groups. The bar had a high frequency of occasional visitors, including people experiencing homelessness and those affected by alcoholism. Despite considerable efforts to stop the outbreak, it continued to spread over a period of years.

**Figure 7:**
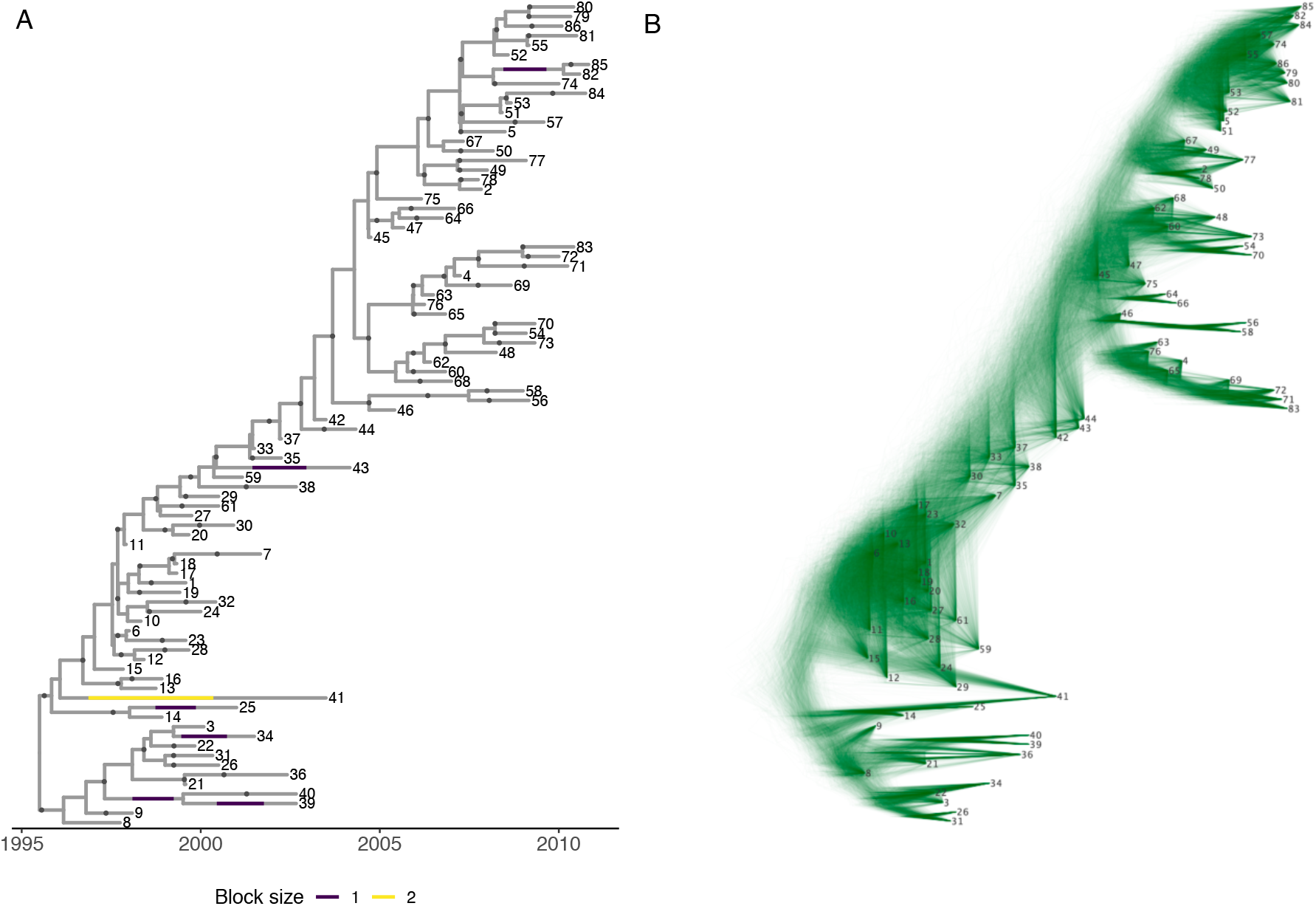
Results from TB outbreak analysis with labels the same as in Figure 6. A. maximum a posteriori (MAP) tree with blocks and transmissions shown. B. DensiTree [4] of the transmission tree analysis, showing considerable uncertainty in both clade support and internal node heights.

TB arises from a process of person-to-person transmission with a variable time frame of years from infection to infecting others and to sampling (as reflected in our intensity functions). Including this information, as BREATH does, informs the phylogenetic analysis and results in a phylogenetic tree requiring less overall evolution to explain the data than a coalescent model. For context, Figure 7 shows the MAP annotated phylogenetic tree from our log, and well the DensiTree plot of our posterior sample. We also compare BREATH’s transmission inferences with those of TransPhylo [17] (see Supplementary Materials). BREATH has more higher-probability direct transmission pairs than TransPhylo at these parameter settings (Figure S3), probably because there are fewer constraints on the transmission trees with multiple phylogenetic trees, whereas TransPhylo has a fixed phylogeny. Commensurate with having more higher-probability direct transmission pairs overall, BREATH also has more higher-probability transmission pairs with a SNP distance larger than 1 or 2, though these are by far in the minority (Figure S4).

## 4. Discussion

BREATH is a Bayesian method that simultaneously constructs phylogenetic and transmission trees, accounting for within-host evolution and allowing for a flexible number of unsampled individuals, and real-time outbreaks. Duault *et al*. reviewed transmission reconstruction methods [30], listing methods according to (1) the data needed (here, sequences and sampling times), (2) whether the methods are phylogenetic or not and (3) how unobserved processes are modelled. BREATH is phylogenetic, not sequential, in their language. BREATH uses intensity functions, which results in a highly flexible model that can incorporate a latent period; it does not use an explicit spatial model, migration model or a susceptible-exposed-infectious-recovered model.

The BREATH tree prior leverages BEAST 2’s power in phylogenetic inference: it can be combined with nucleotide, aminoacid and other types of sequence data, various models of evolution of characters along a tree and different clock models.^4^ BREATH overcomes a substantial barrier in reconstructing transmission histories, since previous methods could not account for unsampled hosts while estimating phylogenetic and transmission trees, and cannot account for preferentially observing faster-occurring events in real-time analyses (the right truncation). We applied the method to data from a TB outbreak [7], and found that our transmission process led to more well-resolved phylogenetic trees than standard models. We identified several distinct stages of transmission with only a few concurrent hosts separating them, suggesting that the outbreak came very near to being interrupted in the early 2000s. While there is considerable uncertainty in who infected whom, we identified some high-probability transmission events, distinct subgroups in the outbreak, and found some hosts likely infected by an individual not in the sample. Information about the posterior numbers of secondary infections for each host, which hosts did not have a plausible infector among the sampled individuals, the posterior times between infection and sampling and similar information can readily be extracted from the posterior transmission trees in our model. These are all potentially useful even with uncertainty in individual transmission events.

BREATH has the limitation that there are multiple interacting parameters that influence branch lengths and therefore also the estimated numbers of unsampled individuals. How these interact, and which part of parameter space enables accurate inference of phylogenies and transmission histories, is still to be determined and will vary by pathogen. As in previous methods, it is not likely to be possible to infer the generation time (from infection to infecting another), the within-host parameter and the numbers of unsampled individuals: without some prior information, there would be no way to distinguish between a branch with many brief inter-transmission intervals or fewer longer ones. For most infectious diseases for which sequence data are likely to be available, knowledge of the typical timing between exposure and symptoms (even if, as with TB, these may be highly variable), the duration of infectiousness and/or the relationship between symptoms and infectiousness mean that there is some available information to inform the time scale and range of the distribution for the time between exposure and infecting others, be it days to 1-2 weeks (COVID-19), weeks (mpox, measles, Ebola virus), 1-3 months (*Streptococcus pneumonaie*) or many months to years (tuberculosis).

There are several changes that could be made to improve the model. For example, the model assumes a host stops infecting others after being sampled, but this is readily modified. It would also be straightforward to relax the assumption of constant within-host *N*_*e*_*g* over time and across hosts. An alternative model may be more appropriate. Different hosts (according to clinical data or risk factors) could also have different within-host parameters, if there were data to inform these. These changes would be straightforward as long as the relevant coalescent likelihood is tractable.

In closely-related outbreaks, the amount of detectable genetic variation is likely to be low, resulting in both phylogenetic uncertainty and uncertainty about who infected whom. For infections for which the sampling time is more informative about infectiousness (than it is for a chronic infection like TB), timing may resolve some of this uncertainty. In any case, it is an advantage of our approach that the phylogeny can be informed by the transmission process, since this is the process that generates the data. Phylogenetic trees are used to understand the acquisition vs transmission of antibiotic resistance in TB [8], to resolve times of origin of emerging clades [33]. and to infer how pathogens move geographically [11]. In any of these applications, having a better phylogenetic model is helpful even if person-to-person transmissions are not the focus of interest.

BREATH assumes that hosts infect other hosts only once, and a single lineage is transmitted. However, hosts could be infected multiple times, depending on immunity, infectiousness and other factors. Knowledge of multiple infections would require multiple samples per host and/or deep sequencing sufficient to characterize individuals’ diverse infections. This raises challenges: the priors or constraints on how a single host’s taxa should be placed in the phylogeny depend on whether the host was infected more than once. If reinfections could be assumed to proceed similarly to other infections, BREATH would be readily adaptable to having multiple samples per host, and each augmented phylogeny would indicate, for that posterior sample, whether the host had one infection that diversified (corresponding to all of that hosts’ taxa contiguously coloured) or whether there was a reinfection. BREATH’s current implementation, while allowing for within-host evolution (different sequences from one host may be transmitted to others, or the sampled sequence from a host may not be the same as the sequence that was transmitted to another), it does not currently account for observed diversity (multiple sampled sequences per host).

The potential benefits of transmission tree inference are not only in human-to-human outbreaks but also in the management of zoonotic infections (such as bovine tuberculosis, foot and mouth disease, avian influenza). Who-infected-whom information is useful in informing policy decisions about whether to quarantine certain areas, cull herds (in case of animal disease), or put movement restrictions in place. In this paper, we modelled hosts as human individuals; hosts could also represent farms affected by a disease outbreak, or non-human hosts. In some contexts there may be information about the transmissibility and/or susceptibility of different groups of hosts, like children being more or less susceptible or infectious than adults. These aspects could be modelled with different intensity functions for different kinds of hosts. This is straightforward in the likelihood for sampled individuals, but unsampled individuals would be challenging to model. More optimistically, integrating contact data by adding priors/constraints on the colouring of a tree would be fairly straightforward.

The transmission tree likelihood is not particularly computationally intensive compared to the sequence tree likelihood, and BEAST 2 readily scales to many hundreds of sequences. Our model can therefore scale well with larger numbers of samples on the scales for which we would anticipate having densely-sampled sequences (see Section S7 for details). Another advantage is that due to the placement of unsampled transmission chains on branches, a preliminary clustering step – identifying putative transmission clusters for onward analysis, with a SNP cutoff or phylogenetic method – should not be required. In the event of an ongoing outbreak, it would be desirable to have a version that allows re-use of previous analyses when new data becomes available (instead of running the whole analysis from scratch.)

## Acknowledgments

This work was funded by the Natural Sciences and Engineering Research Council (Discovery Grant RGPIN-2019-06624) and the Federal Government of Canada’s Canada 150 Research Chair program. We thank Lars Berling for transmission tree extraction and visualization tools, and Jennifer McNichol for a figure contribution.

## A. Supplementary Materials

### S1. Transmission process

#### S1.1. Within-host coalescent likelihood

The second term in the decomposition in (3), *P* (*G*|*T, N*_*e*_*g, τ*_*s*_), is the likelihood of the phylogeny conditional on the transmission tree and the within-host coalescent parameter. As in previous work (BEASTLIER, TransPhylo, phybreak), computing this likelihood is aided by the fact that the transmission tree breaks the phylogeny into smaller, independent subtrees (one for each host), each occurring in a different individual, along with linear chains of transmission which are “trivial” genealogies of just one lineage. Accordingly, the phylogeny likelihood is the product of likelihoods of smaller phylogenies; we use a coalescent model for the within-host evolution.

The within-host parameter *N*_*e*_*g*, intuitively, tells us the mean time that we can expect two lineages to persist before coalescing. For example, in the schematic tree shown in Figure 2, if we do not condition on coalescent events occuring with the infection time, we would have a coalescent likelihood of

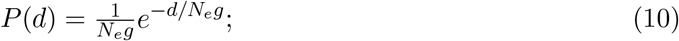

the time to coalescence is exponential with mean *N*_*e*_*g* for two lineages.

Let *G*_*i*_ be a timed genealogy corresponding to a single host, and let *τ*_*i*_ be the times of the leaves for this tree. (The sampling event for a sampled host’s genealogy will be one of the leaf times, and also one of the times *τ*_*s*_. Other tips in host *i*’s genealogy correspond to times at which *i* infected another individual.) Let 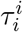 be the time of infection of host *i*. We model a strict bottleneck at transmission, and consequently, all lineages in *G*_*i*_ must coalescence after 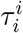 (forward in time), or before 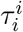 (thinking backward in time). We use the bounded coalescent calculation in Carson *et al* [7] to perform this conditioning. We note, however, that in the event that the infection event for the host is very close to the time of transmission or sampling, such that the bounding time approaches zero, this conditioned likelihood can diverge to infinity. This should be prevented by making very brief host durations *τ* unlikely enough that the combined likelihood does not diverge. While the Carson et al conditioned likelihood has no closed form, and a full exploration of the possible divergence of the bounded coalescent is beyond the scope of this work, choosing the shape parameters *A*^*tr*^ and/or *A*^*s*^ to be large enough should ensure that very short host durations are of sufficiently low likehihood that the overall divergence is avoided.

To see this, consider the example of a two-taxa tree, where the first taxon occurs a time *τ* after the time of infection (forward in time) and the next occurs later. There is therefore a maximum time interval equal to *τ* during which there can be two contemporaneous lineages, and we wish to explore the effect of conditioning on the coalescent event happening within that time. Let *t* be measured backwards in time from time *τ*. The unconditioned likelihood that the two lineages coalesce at time *t* = *T* is 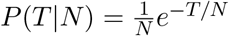. The conditioned probability with *T < τ*, requires dividing by the probability that *T < τ* given *N* :

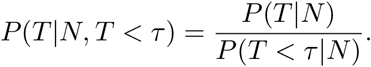

In this simple example, the denominator is given in Wakeley [2] as *g*_2,1_(*T*):

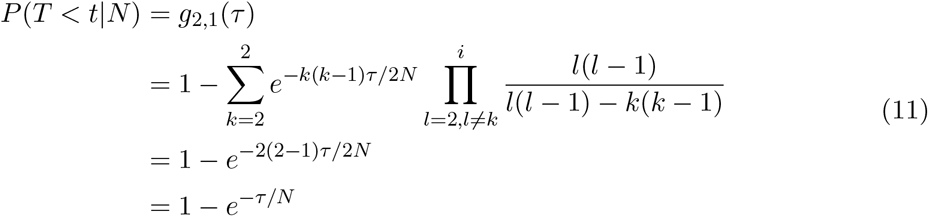

The conditioned probability *P*_*c*_ is then

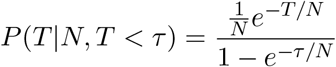

which diverges towards *∞* as *τ →* 0. This is as expected: the distribution should approach a *δ* function as *T* approaches 0. But it can cause numerical issues, and an MCMC might propose very short branch lengths and obtain high likelihoods.

However, we can consider this steep increase in likelihood in combination with the likelihood of *τ* given the parameters guiding the host’s duration of infection and time to sampling. If the tip at time *τ* is the only transmission event from this host, *τ*’s density is described by the gamma distribution with shape *A*^*tr*^ and scale *B*^*tr*^. Consider now the joint likelihood of *τ* and time to coalescence *T*, conditional on *N* and *T < τ* :

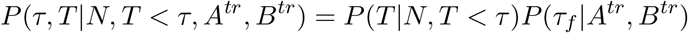

For readability, let *α* = *A*^*tr*^ and *β* = *B*^*tr*^. We have

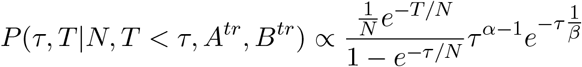

By L’Hopital’s rule, as long as *α >* 2, this goes to 0 as *τ →* 0. While this is only for a two-taxon tree, each additional taxon within the host will correspond to an additional sample from the generation time gamma distribution with parameters *α* and *β* and (loosely) an additional term *τ*^*α−*1^ in the numerator of an expression similar to the one above. A detailed exploration of the behaviour of the joint probabilities of host times (and consequently bounding times) and within-host trees under the bounded coalescent is beyond the scope of the current paper.

BREATH allows the user to choose not to use the bounded (conditioned) coalescent, or to set a minimum allowable host duration.

Intuitively, the parameter *N*_*e*_*g* determines how likely is it for members of the within-host population branches to share a direct ancestor, so it reflects the within-host evolution. When *N*_*e*_*g* is large, the coalescent rate within the host is small, and it takes more time for lineages that are both present to coalesce. Conversely, if *N*_*e*_*g* is small they coalesce much faster, and multiple branches are less likely to be present at the same time. While the version here assumes *N*_*e*_*g* to be constant, other parametric forms can be assumed, for which see [3].

A “trivial” genealogy *G*_*i*_ occurs when a host only ever has one lineage (a line, with no branching events). This does not require or permit any coalescent events. Its probability is 1 in the coalescent model. However, each internal node in the phylogeny is modelled as within some host, either a sampled host or an IUH. The genealogies corresponding to these hosts are not trivial, because they contain one or more branching events.

#### S1.2. Within-host coalescent prior

To obtain a prior for *N*_*e*_*g* we used temporally sampled allele frequency data from a study on the within-host evolution of *M. tuberculosis* [6]. There are methods to estimate *N*_*e*_ in units of generation time (in other words, *N*_*e*_*g* from such data, and we used the framework in Jonas et al [4] (Equations 2 and 3 of that work, for a simple estimate, and a modification of Equation 12, under the assumption that the numbers of bacteria and reads are both large; we do not have this information). We explore using synonymous SNPs only (vs all SNPs), using SNPs not associated with antibiotic resistance only, using all SNPs and using Eqs 2 and 3 vs 12, as well as using different pairs of adjacent temporal samples (0 to 2 weeks, 2 to 4 weeks, and so on). We consistently obtain *N*_*e*_*g* in the range [0, 1]. For a conservative prior, allowing for the fact that TB is haploid, and that this estimate is rough and may be too restrictive, we use a uniform prior of *N*_*e*_*g ∈* [0, 10] for this analysis.

#### S1.3. Derivation of *p*_0_

We obtain *p*_0_, the probability of being unsampled and having all descendants unsampled (i.e. of not being ATTS) with a standard technique from branching processes: conditioning on the number of descendants. Because we will deal with right truncation momentarily, here, we let *p*_0_ be the probability of *ever* being known, and *q* the eventual sampling probability 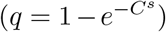:

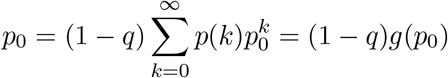

where *g*(*s*) is the probability generating function for the offspring distribution *p*(*k*). In our process, this is Poisson with mean *C*^*tr*^ if sampling does not stop infectors from infecting others after sampling (but for *p*_0_ and related computations in this section, the relevant cases are unsampled). For the Poisson distribution with mean *λ, g*(*s*) = *e*^*λ*(*s−*1)^. We use Newton’s method to compute *p*_0_ numerically. Now we can build the conditioning term 1 *− S*^*E*^(*Y*_*R*_).

#### S1.4. Probability of being once-ATTS and multiply-ATTS

The probability that an individual is not ancestral to the sample (ATTS) is *p*_0_. The probability that an indvidual *j* is “once ATTS” is the probability that precisely one of *j*’s infectees is ATTS,

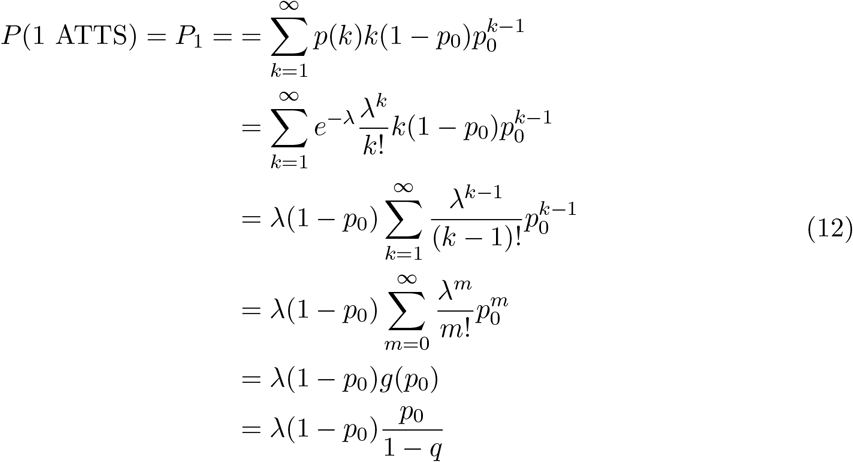

where *g* is the probability generating function for the distribution *p*(*k*), which we model as Poisson, and the final line follows from the form of *p*_0_ in the main text. We use *λ* = *C*^*tr*^.

In order to be what we have called “twice-ATTS”, an individual must simply not be unknown, and not be only once ATTS, so we have

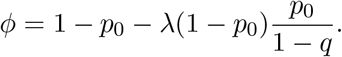

In the case of absolutely perfect, complete, sampling (never any chance of an unsampled case), *p*_0_ = 0 and *ϕ* is not relevant because we would not have chains of unsampled cases. We have not implemented this possibility.

#### S1.5. The likelihood for chains of transmission

Recall that the chain of unsampled transmission ends if either someone is sampled or they infect someone who is MATTS. We use an intensity function *h*^*e*^ to describe this: *h*^*e*^(*t*) = *h*^*s*^ + *h*^*tr*^*ϕ*, where *ϕ* is as defined above. This allows us to find the success probability for the geometric distribution. The probability *ρ* that an event from an intensity function *h*^*e*^ happens is one minus the probability that it never occurs:

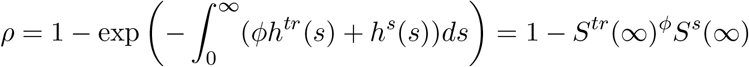

Note that 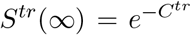 and similarly for *S*^*s*^. We need *∞* in the argument, rather than a right censoring time, because we need to account for right truncation using the general principle *f* (*t*|*t < Y*_*R*_) = *f* (*t*)*/*(1 *− S*(*Y*_*R*_)), as described in the main text, so we need *f* (*t*), the density for the time if there were no truncation, in the numerator. In this case, *Y*_*R*_ is 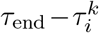 (the infection time of the block). We have *p*(*n, t*) = *p*(*n*)*p*(*t*|*n*) where *p*(*n*) = (1 *− ρ*)^*n*^ and 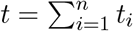 where *t*_*i*_ is the length of time it takes for an individual in the chain to infect the next individual. We model *t*_*i*_ *~* gamma(*a, b*), so that *t ~* gamma(*nA*^*tr*^, *B*^*tr*^) as described in the main text.

As described in the main text, these ingredients (in particular *ρ*) complete the likelihood:

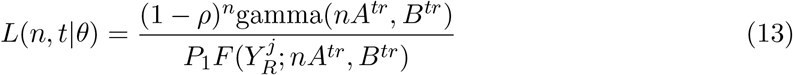

where *n* is the count, 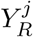 is the block’s right truncation time, *F* is the gamma CDF and *t* is the duration of the block. We have now defined all the ingredients needed to explicitly write, and compute, the transmission likelihood in (4), and in fact, the whole likelihood in (3).

### S2. Hastings Ratios

We have minor changes to the dimensionality when moves change whether a branch has an infection event or not (block count *b*_*c*_ changes between −1 and 0) or whether a branch has a chain of unsampled transmission (*b*_*c*_ *≥* 1) or not. We also define moves in terms of fractions of branches (with parameters in (0, 1)) but the posterior depends on times (in units of branch lengths).

We use the following points repeatedly:

1. Suppose we sample *b*_*s*_, *b*_*e*_ on a branch *b* of length *l*(*b*). Since *b*_*s*_ *< b*_*e*_, the density for (*b*_*s*_, *b*_*e*_) is 2*/l*(*b*)^2^, which is the inverse of the area of the triangle with equal sides of length *l*(*b*).
2. If we sample *b*_*µ*_, the position of the infection event on branch *b* with *b*_*c*_ = 0, the density is 1*/l*(*b*).
3. The HR for a move where each parameter is sampled in (0, 1) and then simply scaled by a length *l*(*b*), and one where it is sampled in (0, *l*(*b*)) directly, are the same. In other words, whether the volume change is handled in the density of sampling a new variable (uniformly), or in the transformation from the unit interval to a branch length, the resulting HR must be the same.
4. Accordingly, in the language of Green [1], we handle the parameter sampling with reference to densities *g*(*u*) and *g*(*u*^*′*^), where the forward move requires sampling one or more parameters *u* and the reverse move *u*^*′*^.

We let |*B*_elig_(*x*)| be the number of branches on which it is possible to remove an infection if the state is *x*. When an edge has *b*_*c*_ = 0, this denotes an infection on the edge which happens at a point *b*_*µ*_ on the edge (the “start” and “end” values for this zero-duration “block” are the same, and we denote this point *b*_*µ*_). First we discuss the block operators, and then the more complicated case of the infection movers.

#### Block operators

1. **Block boundary operator**: we either move *b*_*µ*_, *b*_*s*_ or *b*_*e*_. There is no change of dimension, no new variables, and no change to *B*_elig_. The HR is 1.
2. **Block addition and removal operator**: Consider the moves that remove an infection from branch *b*. There are several variations, depending on *b*’s block count. Each variation will have the same move probability ratio *j*_*m*_(*x*^*′*^)*/j*_*m*_(*x*), which is equal to

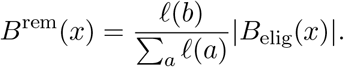
  a. A block *b* with *b*_*c*_ *>* 1 continues to have a block:

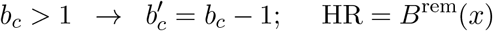
  b. A block *b* with *b*_*c*_ = 1 becomes an “empty block” with *b*_*c*_ = 0, so *b*_*s*_, *b*_*e*_ *→ b*_*µ*_. The new variable in the forward direction is *b*_*µ*_ and in the reverse direction there are two new variables, *b*_*s*_ and *b*_*e*_. This gives

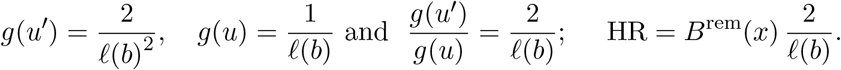
  c. A block *b* with *b*_*c*_ = 0 loses an infection so 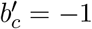. Here,

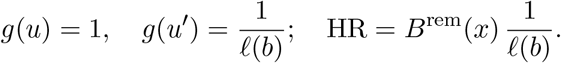
  The reverse moves, adding infections, have the appropriate inverse HRs.

#### Infection movers

The infection movers are more complex, because there is a source branch *b* and a destination branch *d* when we move an infection from one branch to another. This means that we need to account for changes in the block counts and boundaries on the two branches, and the differences in branch lengths.

Recall that the narrow version moves an infection from *b* to *b*’s sibling branch, and the wide version the infection to another branch chosen with probability *l*(*b*)*/ Σ* _*a*_ *l*(*a*). For the narrow version, the move probability ratio is 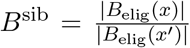 (where *x* denotes the state of the tree before the move and *x*^*′*^ after it).

For the wide version, the probability of the move is the probability that we choose *b* and the destination *d*: 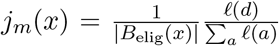. The same applies in reverse: we need the probability that we would start with *d*, remove an infection from it and put it on *b*. This means that the move probability ratio is

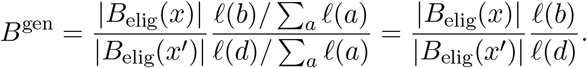

Accounting for the scaling between (0, 1) and the relevant branch lengths, the remaining HR computations reduce to the appropriate ratios of these densities. There are 9 cases, some of which are the reverse of others. In each case, we use *B* to denote the appropriate choice of *B*^sib^ or *B*^gen^. In the table, the transformation and reverse transformation columns refer to the inverse volume of the parameter choice in the forward and reverse moves, respectively.

#### Hastings Ratios for the infection movers

| Case | Source branch | Destination branch | Tranformation | Reverse transformation | HR |
| --- | --- | --- | --- | --- | --- |
| 1 | $b_c = 1, b'_c = 0$<br>$(b_s, b_e) \rightarrow b_\mu$ | $b_c = -1, b'_c = 0$<br>$\{\cdot\} \rightarrow b_\mu$ | $\frac{1}{\ell(b)\ell(d)}$<br>reverse of 6 | $\frac{2}{\ell(b)^2}$ | $2B \frac{\ell(d)}{\ell(b)}$ |
| 2 | $b_c = 1, b'_c = 0$<br>$(b_s, b_e) \rightarrow b_\mu$ | $b_c = 0 \rightarrow b'_c = 1$<br>$b_\mu \rightarrow (b_s, b_e)$ | $\frac{1}{\ell(b)} \frac{2}{\ell(d)^2}$<br>reverse of itself | $\frac{2}{\ell(b)^2} \frac{1}{\ell(d)}$ | $B \frac{\ell(d)}{\ell(b)}$ |
| 3 | $b_c = 1, b'_c = 0$<br>$(b_s, b_e) \rightarrow b_\mu$ | $b_c \geq 1 \rightarrow b'_c \geq 1$<br>$(b_s, b_e) \rightarrow (b_s, b_e)$ | $\frac{1}{\ell(b)}$<br>reverse of 7 | $\frac{2}{\ell(b)^2}$ | $B \frac{2}{\ell(b)}$ |
| 4 | $b_c = 0 \rightarrow b'_c = -1$<br>$b_\mu \rightarrow \{\cdot\}$ | $b_c \geq 1 \rightarrow b'_c \geq 1$<br>$(b_s, b_e) \rightarrow (b_s, b_e)$ | 1<br>reverse of 8 | $\frac{1}{\ell(b)}$ | $B \frac{1}{\ell(b)}$ |
| 5 | $b_c = 0 \rightarrow b'_c = -1$<br>$b_\mu \rightarrow \{\cdot\}$ | $b_c = -1 \rightarrow b'_c = 0$<br>$\{\cdot\} \rightarrow b_\mu$ | $\frac{1}{\ell(d)}$<br>reverse of itself | $\frac{1}{\ell(b)}$ | $B \frac{\ell(d)}{\ell(b)}$ |
| 6 | $b_c = 0 \rightarrow b'_c = -1$<br>$b_\mu \rightarrow \{\cdot\}$ | $b_c = 0 \rightarrow b'_c = 1$<br>$b_\mu \rightarrow (b_s, b_e)$ | $\frac{2}{\ell(b)^2}$<br>reverse of 1 | $\frac{1}{\ell(b)\ell(d)}$ | $B \frac{\ell(d)}{2\ell(b)}$ |
| 7 | $b_c > 1 \rightarrow b'_c \geq 1$<br>$(b_s, b_e) \rightarrow (b_s, b_e)$ | $b_c = 0 \rightarrow b'_c = 1$<br>$\{\cdot\} \rightarrow b_\mu$ | $\frac{2}{\ell(d)^2}$<br>reverse of 3 | $\frac{1}{\ell(d)}$ | $B \frac{\ell(d)}{2}$ |
| 8 | $b_c > 1 \rightarrow b'_c \geq 1$<br>$(b_s, b_e) \rightarrow (b_s, b_e)$ | $b_c = -1 \rightarrow b'_c = 0$<br>$\{\cdot\} \rightarrow b_\mu$ | $\frac{1}{\ell(d)}$<br>reverse of 4 | 1 | $B\ell(d)$ |
| 9 | $b_c > 1 \rightarrow b'_c \geq 1$<br>$(b_s, b_e) \rightarrow (b_s, b_e)$ | $b_c \geq 1 \rightarrow b'_c > 1$<br>$(b_s, b_e) \rightarrow (b_s, b_e)$ | 1<br>reverse of itself | 1 | $B$ |

### S3. Simulator

#### S3.1. Simulator description

We also implemented a simulator independent of the transmission likelihood, that allows us to simulate transmission trees with phylogenies, allowing us to test the model. The inputs for the simulator are the parameters for the sampling and transmission intensity functions (3 each), *N*_*e*_*g* and the stopping time *τ*_end_.

Initialize a list *L* of hosts whose infections are to be simulated. *L* can contain the times of infection, and the infector. Start with one host to be simulated, starting at time *t* = 0. *L* starts out being a row (0, NA). (NA for the fact that this is the index host, and they have no infector in the process).

Simulate an individual host, *j*, with time of infection 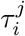:

1. Simulate *n*_*j*_ *~* Poisson(*C*^*tr*^). This is the number of new infections that the host would eventually cause (without considering the stopping time).
2. Simulate whether the host will be sampled: *s*_*j*_ *~* Bernoulli 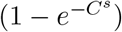. If *s*_*j*_ = 1 the host would be sampled, if the sampling time occurs early enough.
3. If *s*_*j*_ = 1, simulate the time of sampling:

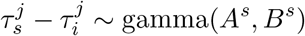

If 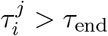, the host is not sampled after all. Ignore 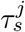.
4. Simulate the times when *j* infects the *n*_*j*_ new infectees:

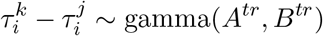

with *k* = 1, …, *n*. Remove any 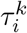 that are above 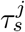 (model A, where sampling stops people from transmitting) or *τ*_end_ (model B, where transmission ends when we end observation).
5. Sampling a within-host phylogeny (see below) for host *i* using the times 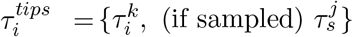 Attach the origin of this phylogeny to the infector of host *j*, accounting for the fact that the tMRCA of this phylogeny is *after* (forwards in time) 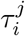, not equal to 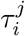.
6. Remove the (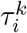, infector for *i*) row from *L*. Append the set 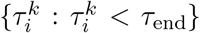 times of infection and *j* as the infector to the list *L* of infection times and infectors. *L* gets rows 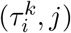 appended to it, only for those 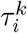 that are less than *τ*_end_.

Continue by simulating the hosts in the list, gluing the phylogenies together. Stop when there are no more hosts to infect. Depending on the intensity function parameters, trees with zero or one taxon often have a high probability, which can be interpreted as an outbreak immediately being stopped. Another mode can often be observed with just a few taxa, indicating the outbreak dies out very early on. But there tends to be a long tail in the taxon count distribution, some reaching a very high number of taxa. If a fixed number of taxa is desired, the tree is rejected if the number of taxa differs from the desired one, and a new tree is generated until one is encountered with the desired number of taxa. Due to the thinness of the tail and depending on the intensity functions’ parameters, this can take considerable time. This also has the potential to produce bias in some tests of the model, because the desired number of taxa may be unlikely given the epidemiological parameters. Our simulation tests do not seem affected by the choice of the number of taxa.

Some of the within-host phylogenies will be trivial. These are sampled hosts who infect no others, or unsampled hosts who infect precisely one other, who is not sampled. These latter become the chains of unsampled transmission. Any host (sampled or not) with two or more tips in 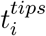 contains at least one node of the larger phylogeny, because it has at least one coalescent event in it.

Figure S1 shows a simulated tree including within host coalescent events and unsampled hosts. The red tree is the one output by the simulator.

The transmission tree simulator is available as the `TransmissionTreeSimulator` app in the `BREATH` package for BEAST 2. It has the following options:

- endTime (real number): end time of the study
- popSize (real number): population size governing the coalescent process
- sampleShape (real number): shape parameter of the sampling intensity function
- sampleRate (real number): rate parameter of the sampling intensity function
- sampleConstant (real number): constant multiplier of the sampling intensity function
- transmissionShape (real number): shape parameter of the transmission intensity function
- transmissionRate (real number): rate parameter of the transmission intensity function
- transmissionConstant (real number): constant multiplier of the transmission intensity function
- out (file name): output file. Print to stdout if not specified (optional)
- trace (file name): trace output file with end time, tree heights and tree lengths, or stdout if not specified
- seed (long): random number seed used to initialise the random number generator (optional)
- maxAttempts (integer): maximum number of attempts to generate coalescent sub-trees (default: 1000)
- taxonCount (integer): generate tree with taxonCount number of taxa. Ignored if negative (default: −1)
- maxTaxonCount (integer): reject any tree with more than this number of taxa. Ignored if negative (default: −1)
- treeCount (integer): generate treeCount number of trees (default: 1)
- directOnly (true|false): consider direct infections only, if false block counts are ignored (default: true)
- quiet (true|false): suppress some screen output (default: false)

**Figure S1:**
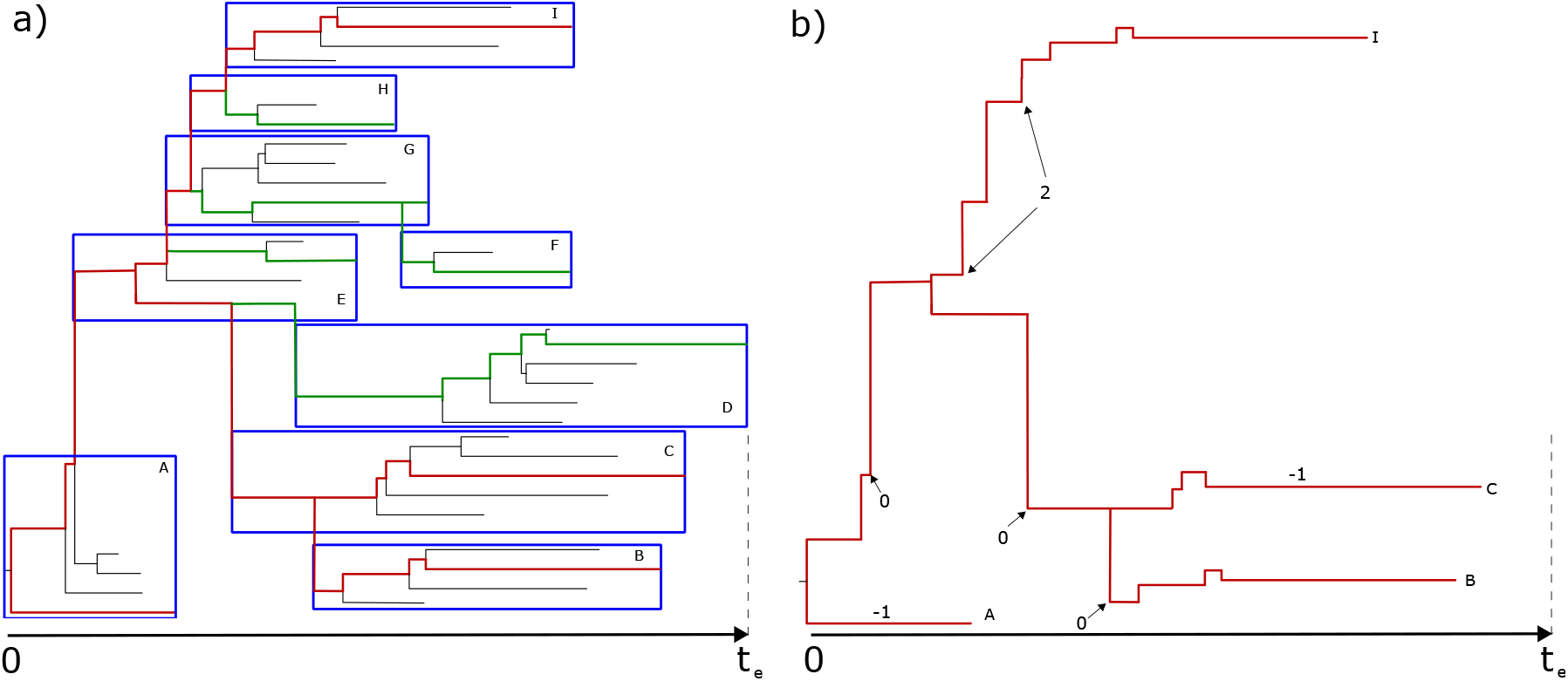
Simulated tree from time 0 to time *t*_*e*_ indicated on the x-axis. a) Small simulated tree where blue boxes indicate hosts, red tree the tree ending in samples, red+green branches are branches generated by the simulator as within host coalescent trees, red+green+black branches form the underlying phylogeny. b) Tree output by the simulator. Numbers on branches are block counts, arrows indicate block start and end of blocks (if block count at least zero). For blocks with count of zero start and end of block coincide. Hosts D to H are not sampled, so these are removed from the simulator output. Host E becomes an unsampled host infected by A and infecting C. Hosts G and H form a block of size 2 of unknown hosts, while hosts D and F leave no trace and remain unknown unknowns.

To use the command line version of the simulator, use the ‘applauncher’ application (which is part of the BEAST 2 distribution) from a terminal/command prompt. Any of the above options can be used.

Alternatively, start BEAUti (which is also part of the BEAST 2 distribution), select the ‘File/Launch apps’ menu, and select ‘TransmissionTreeSimulator’ from the list of applications. Click the ‘launch’ button to start a GUI version of the simulator.

There is a `WIWVisualiser` tool available in the `BREATH` package to create SVG files to visualise who infected whom, based on a posterior sample. The `WIWVisualiser` has the following options:

- trees (file name): tree file file with transmission trees (optional).
- log (file name): trace file containing infectorOf log. Ignored if tree file is specified (optional)
- burnin (integer): percentage of trees to used as burn-in (and will be ignored) (optional, default: 10)
- out (file name): output file, or stdout if not specified (optional, default: /tmp/wiw.svg)
- prefix (string): prefix of infectorOf entry, e.g., infectorOf (optional, default: infectorOf)
- threshold (real number): probability threshold below which edges will be ignored. (optional, default: 0.1)
- partition (string): name of the partition appended to ‘blockcount, blockend and blockstart’ (optional)

There is also an interteractive visualization app at https://github.com/Lars-B/interactive-wiw. To extract transmission trees as data frames listing who infected whom and when, including individual unsampled hosts and blocks, there is a `breath-helper` tool in the `pyccd` package by Lars Berling at https://github.com/Lars-B/pyccd.

#### S3.2. Simulation parameter guidelines

- Choose *C*^*tr*^ *∈* [1, 4]. This sets the scale of the tree.
- Choose the “sample constant” *q ∈* (0.5, 1) and compute *C*^*s*^ = *−* log(1 *− q*). The range is such that we sample enough cases that person-to-person transmission inference is likely to be a reasonable task.
- Choose 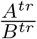 to set the mean inter-infection time (ignoring sampling).
- Choose 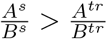 so that sampling occurs at after the mean generation time, on average. Otherwise it seems likely that the transmission chains will die out quickly.
- Choose *τ*_end_, the stopping time. Keep in mind that if the mean time to sampling is considerably greater than the mean time to infection, and *C*^*tr*^ is high, the number of infections could grow very large.
- Choose *N*_*e*_*g* in such a way that the probability that lineages will coalesce in the required time is reasonably high, for example 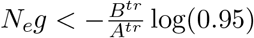.

After choosing the hazard function parameters, a quick sanity check is to plot the gamma distribution densities of the sampling and transmission hazard in the same plot. This plot shows how likely it is for a transmission to happen at a given time and how likely it is for a host to be sampled. For an exponentially growing process, the mean of the sampling hazard should be larger than that of the transmission hazard, especially if sampling prevents subsequent transmission.

Reducing *C*^*tr*^ will make a big (nonlinear) difference. Changing *C*^*s*^ will not make much difference to transmission or the size of the process (though it will to the number of sampled cases, in a linear way), because if sampling happens, it’s most likely to happen after the peak in transmission anyway.

#### S3.3. Simulator troubleshooting

1. Do the transmission chains not take off? e.g. there are no more cases to simulate, but the max sampling times are much less than *t*_*e*_? Suggestion: increase 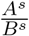 (delay sampling until longer after transmission), increase *C*^*tr*^, or if the transmission process is taking off but there are too few generations (too little time), increase *τ*_end_.
2. Is the number of cases exploding exponentially and there are too many? See above, but do the opposite.
3. Within-host coalescent trees are rejected because they don’t coalesce in time? Decrease *N*_*e*_*g*.
4. Within-host coalescent trees have very very short branch lengths? (This could be OK.) Increase *N*_*e*_*g*.

### S4. Simulation study with varying hazard parameters

**Table S1:**
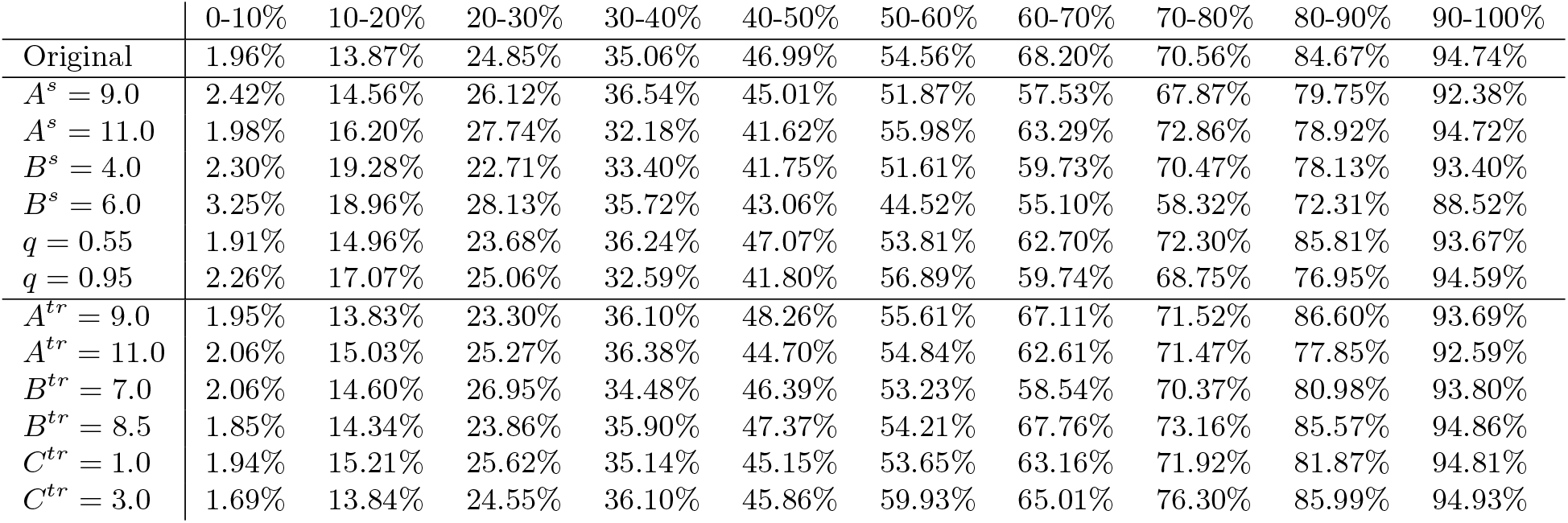
Who-infected-whom inferred probabilities versus true values as in Fig.4 but for a simulation study with varying hazard parameters. The row marked ‘Original’ was run with the same parameters as used to obtain the ground truth, all others have one parameter value changed.

|  | 0-10% | 10-20% | 20-30% | 30-40% | 40-50% | 50-60% | 60-70% | 70-80% | 80-90% | 90-100% |
| --- | --- | --- | --- | --- | --- | --- | --- | --- | --- | --- |
| Original | 1.96% | 13.87% | 24.85% | 35.06% | 46.99% | 54.56% | 68.20% | 70.56% | 84.67% | 94.74% |
| $A^s = 9.0$ | 2.42% | 14.56% | 26.12% | 36.54% | 45.01% | 51.87% | 57.53% | 67.87% | 79.75% | 92.38% |
| $A^s = 11.0$ | 1.98% | 16.20% | 27.74% | 32.18% | 41.62% | 55.98% | 63.29% | 72.86% | 78.92% | 94.72% |
| $B^s = 4.0$ | 2.30% | 19.28% | 22.71% | 33.40% | 41.75% | 51.61% | 59.73% | 70.47% | 78.13% | 93.40% |
| $B^s = 6.0$ | 3.25% | 18.96% | 28.13% | 35.72% | 43.06% | 44.52% | 55.10% | 58.32% | 72.31% | 88.52% |
| $q = 0.55$ | 1.91% | 14.96% | 23.68% | 36.24% | 47.07% | 53.81% | 62.70% | 72.30% | 85.81% | 93.67% |
| $q = 0.95$ | 2.26% | 17.07% | 25.06% | 32.59% | 41.80% | 56.89% | 59.74% | 68.75% | 76.95% | 94.59% |
| $A^{tr} = 9.0$ | 1.95% | 13.83% | 23.30% | 36.10% | 48.26% | 55.61% | 67.11% | 71.52% | 86.60% | 93.69% |
| $A^{tr} = 11.0$ | 2.06% | 15.03% | 25.27% | 36.38% | 44.70% | 54.84% | 62.61% | 71.47% | 77.85% | 92.59% |
| $B^{tr} = 7.0$ | 2.06% | 14.60% | 26.95% | 34.48% | 46.39% | 53.23% | 58.54% | 70.37% | 80.98% | 93.80% |
| $B^{tr} = 8.5$ | 1.85% | 14.34% | 23.86% | 35.90% | 47.37% | 54.21% | 67.76% | 73.16% | 85.57% | 94.86% |
| $C^{tr} = 1.0$ | 1.94% | 15.21% | 25.62% | 35.14% | 45.15% | 53.65% | 63.16% | 71.92% | 81.87% | 94.81% |
| $C^{tr} = 3.0$ | 1.69% | 13.84% | 24.55% | 36.10% | 45.86% | 59.93% | 65.01% | 76.30% | 85.99% | 94.93% |

### S5. Simulation study under SARS-CoV-2 Omicron settings

The simulation studies described in Section 3.1 were done with hazards parameterised with time in years, relevant for tuberculosis. Here, we repeat the simulation study with settings more realistic for an outbreak of SARS-CoV-2. For omicron, the incubation period was estimated [8] to be 3.2 days. The generation time was estimated to be approximately 3 days, but this is the realised generation time, which may be shorter than the intrinsic generation time [9]. The appropriate times would likely be somewhat longer for Delta and earlier variants. We set up simulation model for an Omicron-like outbreak, with a mean generation time of 3.5 days and a standard deviation of 1.5 days, which implies *A*^*tr*^ = 5.4, *B*^*tr*^ = 1.5. We set *C*^*tr*^ = 1.5 implying on average 1.5 infection per host. Then, suppose testing happens after symptoms and severity arise, and organizing and getting a test takes 1-2 days. Let the mean time from infection to testing therefore be 5 days, with a standard deviation of 1 day. This implies *A*^*s*^ = 25 and *B*^*s*^ = 5 and we assume *q* = 0.7, modelling a well-sampled outbreaks. We use a study time of 70 and 100 days, and condition on outbreaks of 32 samples. As before, 100 trees were simulated using the simulator and inferred under the BREATH model using BEAST2. Ground truth trees were also simulated using MCMC (i.e. sampling from the prior), for comparison. We obtained the following coverage and who-infected-whom probability graphs as described in Fig 4:

As with the TB parameter settings, we see a slight deviation between the BREATH inferences and the simulator, with BREATH slightly over-confident on events with high posterior probability, but overall there is a good agreement between the two. The MCMC-based inferences agree even better.

**Table S2:**
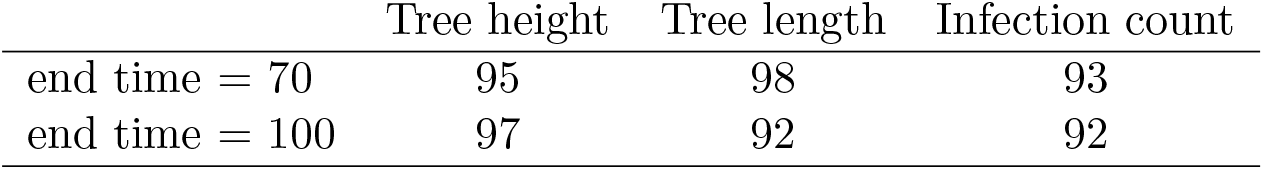
Coverage of the tree height, length and infection count under SARS-CoV-2 parameters.

**Figure S2:**
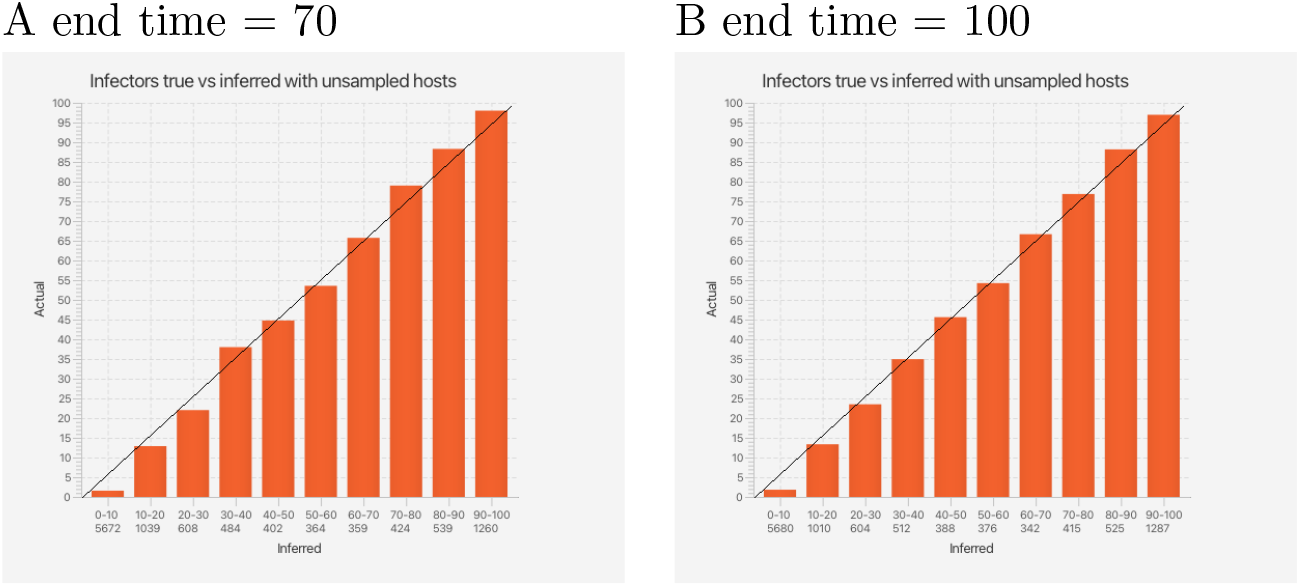
Who infected whom bar plots for the SARS-CoV-2 simulation study

Files used for this simulation study are available from https://github.com/rbouckaert/BREATH/releases/download/v0.0.5/wcss-sars.tgz for the simulator-based analysis, and for the MCMC-based analysis. There is one set of files for the end time of 70, one set for the end time of 100, and one set for MCMC-based ground truth.

### S6. Comparison to TransPhylo

We ran TransPhylo using the same input phylogeny as was used in the original TransPhylo paper [5]. We note that in TransPhylo, if the date of last possible sampling is set to the last tip date, the model may place many unsampled cases near the tips of the tree. This is because of the truncation approach, which is similar to the right truncation used here. Conditioning on the times being less than *τ*_end_ *and* setting *τ*_end_ to equal the last sampling time is conditioning on a very unlikely event, namely: at the last moment in which we could have observed an event, we did. Choosing the bound for the truncation to equal the last time of sampling essentially contradicts the assumptions in the derivation of the decomposition. Accordingly, in our Inference Guidelines (Section S8) and in Table 1 we recommend avoiding this.

We used the following command, i.e. 100,000 iterations, updating the sampling probability, offspring distribution parameter *r* but not *p* and with a date of last possible sampling set to 2015:

~~~
roetztree=read.tree(“roetz.nwk”)
neg=0.03;
recnew=inferTTree(ptreeFromPhylo(roetztree,dateLastSample = 2010.91),w.shape=1.7,
                  w.scale=1/0.3, ws.shape=1, ws.scale=0.5,
                  mcmcIterations=100000,thinning=30, startNeg=neg,startPi=0.5,
                  startOff.r=1,updateOff.r=T,updatePi=T,updateNeg=T,updateOff.p=F,
                  dateT =2015, updateTTree = T,optiStart=T)
~~~

We checked parameter traces for convergence, and extracted who infected whom information with the ‘computeMatWIW’ function with a 20% burnin. TransPhylo’s fixed phylogeny places constraints on the transmission tree, which of course are not constraints in BREATH, because BREATH can change the phylogeny. Figure S3 shows the empirical cumulative distribution functions. Both methods, naturally, have a large number of small-probability events, which is due to the constraint of a transmission tree: there are 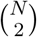 possible pairs and each individual has at most one infector among the sampled individuals, so there are at most *N* − 1 transmission events among sampled individuals in each posterior tree.

**Figure S3:**
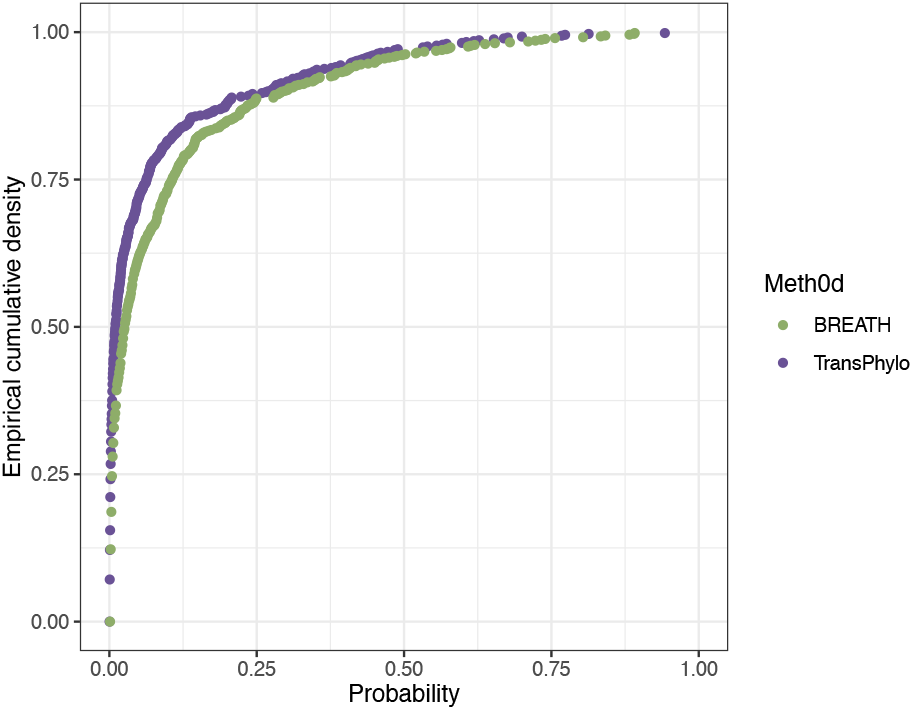
Empirical cumulative distribution functions for the posterior probabilities that *I* infected *j* for all sampled pairs of hosts *i, j* (with *i ≠ j*).

Figure S4 shows the posterior probabilities for sampled pairs and the genetic distances between the pairs. Both methods have higher overall probabilities for pairs whose TB sequences are more similar (low distance), but BREATH shows a more clear pattern of increasing transmission probability for lower SNP distances. Figure S5 compares the posterior probabilities for the highest-probability sources for each sampled host in BREATH with the posterior probability for that same source in TransPhylo. There are some high-probability pairs in BREATH that never appear in TransPhylo, likely because they contradict the fixed input phylogeny. The converse does not occur: all high-probability paris in TransPhylo are recovered by BREATH.

**Figure S4:**
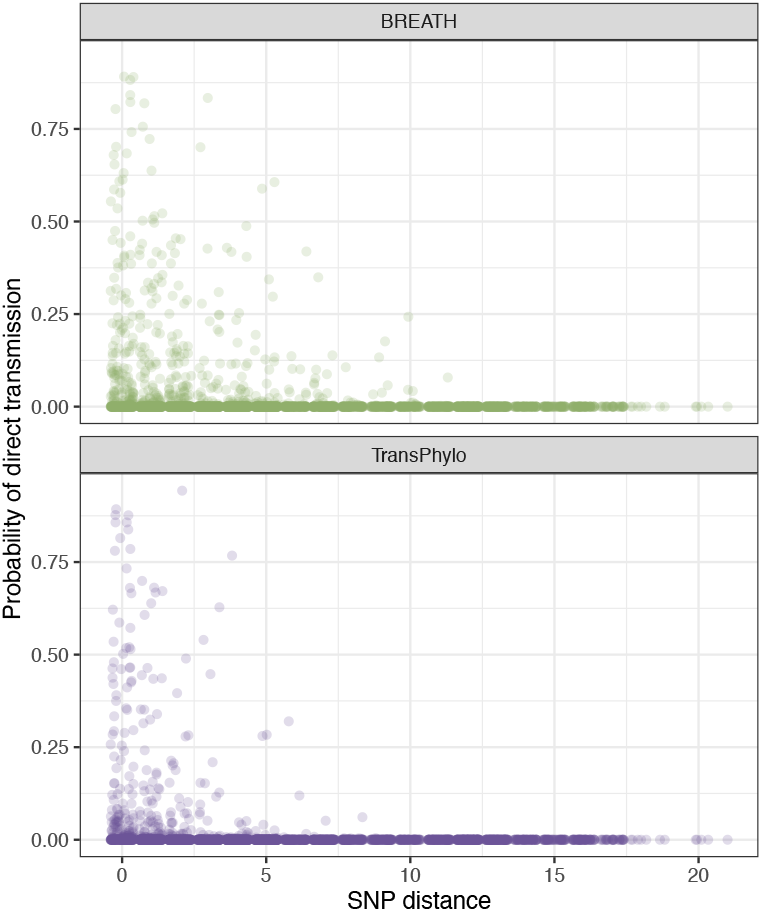
Posterior probabilities for pairs of sampled individuals (the sum of both source-recipient directions for the pair) vs single-nucleotide polymorphism (SNP) distances between the individuals’ TB genomes in BREATH (top) and TransPhylo (bottom).

**Figure S5:**
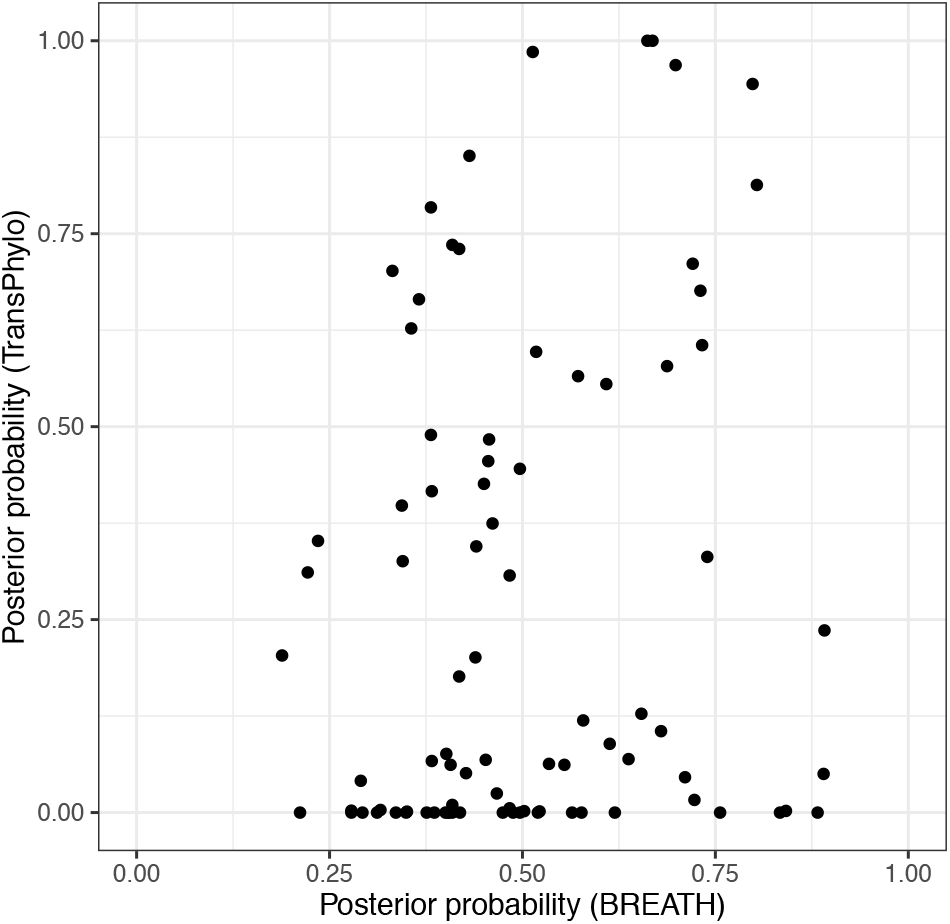
Scatter plot comparing posterior probabilities for the highest-probability source for each sampled recipient in BREATH (x-axis) to the probability of the same source for that recipient from TransPhylo, regardless of whether it was the highest-probability source in TransPhylo. Unsampled sources are included.

### S7. Computational considerations

Giving a general runtime prediction for phylogenetic MCMC is almost impossible: the number of samples required for the MCMC to converge depends on how informative the sequence data is (more informative data requires fewer MCMC samples), how many taxa are involved (up to a few thousand can be done with some perseverance) and how complex other parts of the model is (relaxed clock requires more samples than strict clock), etc.

The analyses for this paper were fast enough that additionally optimising the code was not required, but profiling showed that for the Roetzer analysis about 30% of the time was spent in calculating cumulative gamma probability densities – a linear approximation increased the speed of the MCMC by 25% while introducing only a small error. This suggests the BREATH tree prior can take considerable computational time, where for most analyses the computational time for a single MCMC sample is dominated by Felsenstein’s tree likelihood calculation (usually *>* 95% of the time), and the tree prior only takes a very small amount of time. On the other hand, the Roetzer data consist of a very small alignment (just 85 nucleotides long), which makes it somewhat atypical for phylogenetics, despite it being a somewhat large outbreak given that it is densely sampled at the person-to-person scale. The BREATH calculation is linear in the number of branches, and for each branch linear in the block count on the branch, so for large block counts its calculation slows down. However, the graph in Figure S6 (done on a 2021 Macbook Pro) shows that even with block count 100 per branch, BREATH’s calculation time will be moderate, and is not expected to hamper large analyses, especially when using piecewise linear approximation of the cumulative gamma distribution.

**Figure S6:**
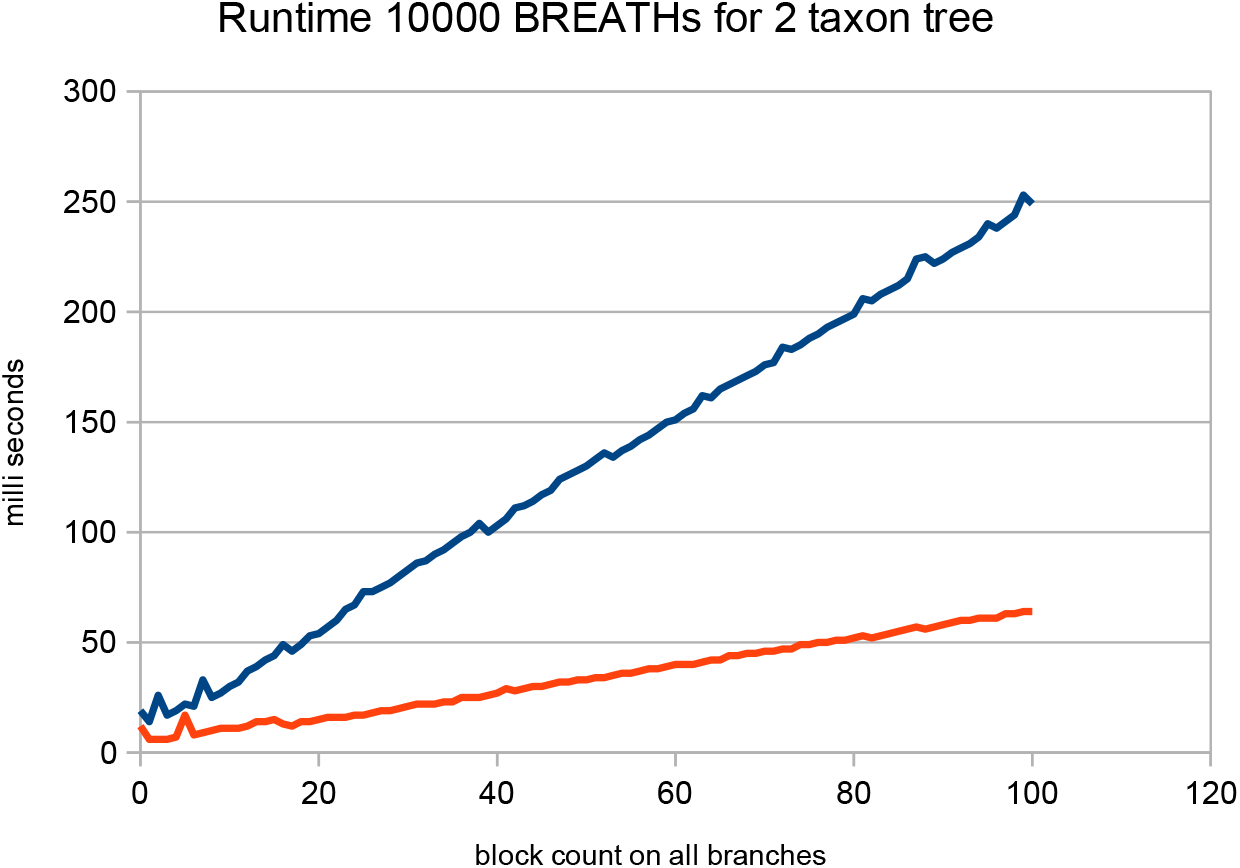
Runtime with exact and piecewise linear approximation for 2 taxon trees. All times remain in the order of milliseconds, making it feasable to use for MCMC. The red line shows the time under the piecewise linear approximation of the cumulative gamma distribution.

### S8. Inference guidelines

Here we provide some guidelines for inferences, and recommendations for users considering whether BREATH is the appropriate model for their data and if so, how to set up the inference.

#### Sampling fraction

BREATH reconstructs person-to-person transmission events, which (generically, not just in BREATH) relies on having sequence data for a high enough fraction of the individuals whose infections are ancestral to the sample for this to be worthwhile. Suppose that overall, in a jurisdiction, there is sequence data for 15% of the incident cases of an infectious disease. With no additional information, this is unlikely to represent enough direct transmission pairs to make person-to-person transmission inference worthwhile beyond simply identifying the most closely-related pairs of sequences and using epidemiological data to determine whether they are likely to reflect direct transmission events. This is because with 15% uniform random sampling, only 2.25% of the *pairs* of individuals are *both* in the sample. However, if a dataset includes one or more densely-sampled transmission clusters, BREATH may be an appropriate tool for transmission and phylogenetic inference *within those clusters* (even where the overall fraction of cases in the whole jurisdiction may be very low). BREATH may also be appropriate for datasets where there are several closely-related sets of sequences to be analysed together, but these are separated by longer phylogenetic branches. Here, BREATH should place chains of unsampled hosts on the long branches and still do a reasonable job at the within-cluster transmission trees. In such a case, the overall sampling could be low, but since unsampled cases are primarily on the long branches separating well-sampled clusters, performance could still be good. Accordingly, there is no fixed rule for how many unsampled cases BREATH can tolerate.

#### Phylogenetic uncertainty

Typically in a person-to-person outbreak, pathogen sequences are closely related. BREATH is suitable for datasets where there is genetic diversity among pathogen sequences and sequences are likely to be informative about transmission, but where there is still phylogenetic uncertainty. If the sequences are very uninformative, BREATH will take most of its information from the hazard parameters and sampling times, and in that case, models based only on symptom times (not included here), sampling times, and epidemiological data may be preferred. If a very well-supported timed phylogenetic tree can be constructed, TransPhylo is likely to be a good approach, but BREATH will accommodate what phylogenetic uncertainty remains.

#### Hazard parameters

BREATH requires knowledge of the hazard parameters and a prior for *N*_*e*_*g* (see Section S3.2). BREATH’s who-infected-whom probabilities are relatively robust to some misspecification of these, but the timing of transmission events may not be as robust, and both caution and sensitivity analysis should be used where the generation time distribution and time-to-sampling distributions are not known. This is particularly the case if the question of interest is how long it takes for individuals to infect others, or to be sampled.

#### Choice of *τ*_end_

The final *possible* sampling time *τ*_end_ should **not** be chosen to be at or very close to the time of the last sample, as discussed above in Section S6. This will result in BREATH (and TransPhylo) placing many unsampled cases near *τ*_end_, because it breaks the assumptions behind the central decomposition, by choosing a prior directly from the data. This results in the model conditioning on an extremely unlikely event. If *τ*_end_ is not known, *τ*_end_ should be set to some considerable time well after max(*τ*_*s*_) on the relevant time scale (i.e. multiple generation times afterwards, or the date at the time of analysis).

#### Choice of *N*_*e*_*g*

The parameter *N*_*e*_*g* is the product of the *within-host* effective population size and the generation time. Neither of these is likely to be known, and both are likely to differ from the *between-host* values. *N*_*e*_*g* may sound obscure, but it matters for the inference, because it affects how long contemporaneous lineages of the tree can be within an individual host, and therefore, it affects how many unsampled cases are needed to “paint” a phylogeny with a transmission tree.

This means that there is a trade-off between *N*_*e*_*g* and the number of unsampled cases, which also has an interplay with the generation time (i.e. the parameters of the transmission hazard), because if it takes longer from infection to infecting another, time elapses (and therefore more of a branch’s length stays) within the same host. Consider the two-taxon “cherry” phylogeny in Figure S7, corresponding to a single transmission event. Without additional sequences, a transmission pair in which *A* infects *B* is always a cherry like this one in our model. The time when two lineages coexist is *d*, and *E*(*d*) = *N*_*e*_*g* (neglecting for the moment the conditioning on coalescence before *A*’s infection time). We must have *d < τ* ^*A*^ and *d < τ*_*g*_, where these times are *A*’s time to sampling and time to infecting B, respectively. It is not coherent to choose the parameters of the model such that the expected time *d* is typically or often larger than the times *τ*_*g*_ and *τ*_*s*_ (set by the transmission and sampling hazards, respectively), because transmission pairs do not make much sense under such settings. If *N*_*e*_*g* is chosen such that *E*(*d*) is much, much smaller than typical values of *τ*_*s*_ and *τ*_*g*_, we can expect large number of unsampled cases to be added. If the known sampling fraction is high, this is contradictory information. So if the sampling fraction is high, *N*_*e*_*g* must be not too different from the time scale of *τ*_*s*_ and *τ*_*g*_ in order for transmission events and their timing to make sense. There are other observations from this small example: 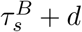 is just under 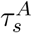 (and the difference is the length of the ‘root’ branch). *d* must therefore be at most the difference in *A*^*′*^*s* and *B*^*′*^*s* time to sampling (where *A*^*′*^*s* is longer, otherwise the cherry would have *B* infecting *A*. While it is not possible to construct hard-and-fast rules for the joint parameter settings, *N*_*e*_*g* and the hazard parameters should be chosen such that an individual transmission pair is coherent on a two-taxon phylogeny, and Figure S7 illustrates how this intuition is developed.

**Figure S7:**
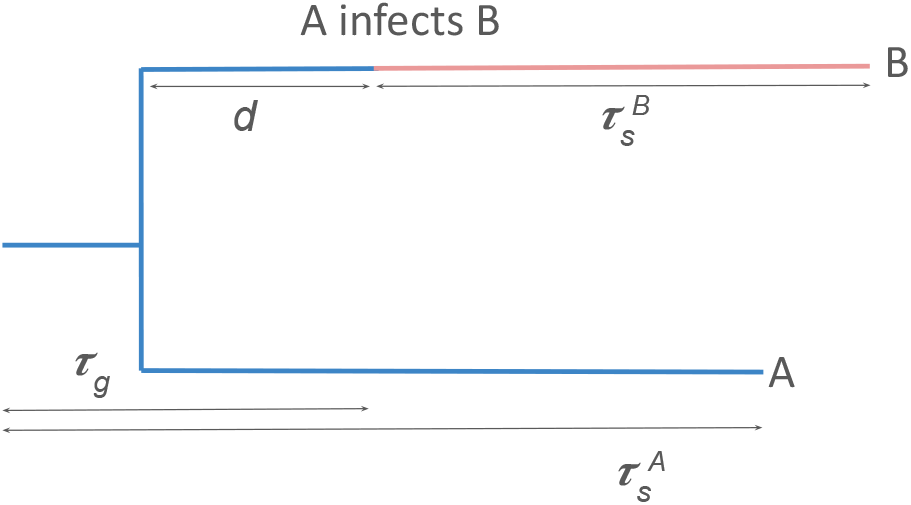
Two individuals on a phylogeny with two taxa

## Footnotes

1 See https://github.com/rbouckaert/BREATH/blob/main/src/breath/operator/TreeWrapOperator.java for an implementation of the tree wrap operator.

2 Files for the well calibrated simulation study are available from https://github.com/rbouckaert/BREATH/releases/tag/v0.0.5.

3 BEAST 2 XML files for the TB analysis are available from https://github.com/rbouckaert/BREATH/releases/tag/v0.0.5.

4 A tutorial for setting up an analysis with BREATH using BEAUti – the user friendly graphical interface for BEAST 2 – is available at https://github.com/rbouckaert/BREATH/tree/main/doc/tutorial.

## Notes

https://github.com/rbouckaert/transmission

### Competing Interest Statement

The authors have declared no competing interest.

### Summary of Updates

Additional explantory figures have been added, and the method has been improved in a number of ways (related to the conditioned coalescent, the MCMC moves and their Hastings ratios). There is a table of notation and expanded simulation studies.

